# Pervasive binding of the stem cell transcription factor SALL4 shapes the chromatin landscape

**DOI:** 10.1101/2025.11.14.688441

**Authors:** Kashyap Chhatbar, Sara Giuliani, Timo Quante, Beatrice Alexander-Howden, Jim Selfridge, Jacky Guy, Tatsiana Auchynnikava, Christos Spanos, Tricia Mathieson, Guido Sanguinetti, Adrian Bird, Raphaël Pantier

## Abstract

Mechanistic understanding of how gene activity is regulated has focussed on the roles of transcription factors at promoters and enhancers, whereas mechanisms capable of globally fine-tuning gene expression through dispersed binding across large genomic regions have received less attention. Here we provide evidence that the essential stem cell transcription factor SALL4 modulates gene expression according to DNA base composition by reading the frequency of its AT-rich target motifs. Using an acute depletion strategy, we establish that SALL4-repressed genes localise to AT-rich genomic domains with high levels of dispersed SALL4 occupancy. While SALL4 is localised within peaks and distributed broadly across the genome, explainable machine learning revealed that its occupancy across the gene body is a strong predictor of transcriptional output. We observed rapid increases in chromatin accessibility and histone acetylation independent of transcriptional activity, suggesting that SALL4 primarily acts upon chromatin, while transcriptional changes are secondary. Accordingly, preventing SALL4 from recruiting the histone deacetylase and nucleosome remodelling corepressor NuRD mimicked a *Sall4*-null phenotype in stem cells and animal models. Our findings reveal that SALL4’s interpretation of DNA sequence optimises the global epigenome and transcriptome, a process integral to maintaining the stem cell gene expression programme.

## Introduction

Mammalian chromosomes are divided into large segments with distinct nucleotide composition, reflecting the non-random distribution of A/T and G/C nucleotides along mammalian genomes ^1,2^.These base compositional domains are evolutionarily conserved ^3,4^ and correlate with structural features including lamina-associated domains ^5^, early/late-replicating regions ^6^ and topologically associating domains ^7^. However, whether AT-rich and GC-rich domains are themselves a response to evolutionary selection, or are a passive consequence of other aspects of genome evolution remains a matter of debate ^4,8^. The importance of GC-rich domains is well illustrated by CpG islands, which coincide with most mammalian gene promoters and interact with several GC-binding proteins ^9,10^, but the biological relevance of AT-rich domains is less well understood. In a search for novel “readers” of DNA base composition, we previously hypothesised that proteins recognising short and frequent motifs made up of only A/T or only G/C base pairs would be able to discriminate fluctuations in nucleotide content across the genome ^11^. Following a proteomic screen in mouse embryonic stem cells (ESCs), we identified the pluripotency factor SALL4 – a member of the Spalt-like family of multi-zinc finger transcription factors ^12^ – as a chromatin protein that can bind a broad spectrum of short A/T-rich motifs ^13^.

SALL4 is highly expressed in stem cells, where it stabilises the pluripotent state, and is indispensable for embryonic development ^14,15^. In humans, SALL4 haploinsufficiency is responsible for a rare congenital disorder called Okihiro syndrome ^16,17^, while its overexpression is commonly observed in a wide range of cancers with poor prognosis ^18,19^. Despite its important physiological and biomedical relevance, relatively little has been known about the molecular basis of SALL4 function. Recently, however, it was reported that SALL4 binds to AT-rich motifs in duplex DNA via its C-terminal C2H2 zinc-finger cluster, ZFC4 ^13,20–22^. Mutations in ZFC4 that abolish DNA binding result in embryonic lethality in mice ^13^ and cause Okihiro syndrome in human patients ^21,23,24^, thereby phenocopying loss-of-function of the entire protein. Transcriptomic analysis showed that mutations in ZFC4 led to preferential up-regulation of AT-rich genes associated with cellular differentiation ^13^. While these findings indicate that sensing of AT-rich DNA base composition by SALL4 is critical for stem cell maintenance and embryonic development, the relationship between SALL4 genomic binding and transcriptional output remained unclear. Whereas *in vitro* assays predicted promiscuous binding to AT-rich motifs dispersed across the genome ^13,20–22,25^, ChIP-seq analysis in pluripotent cells detected SALL4 enrichment at enhancer elements marked by H3K27ac/H3K4me1 ^13,26,27^. Localisation of SALL4 at “active” regulatory elements was not easily reconciled with the high frequency of its (A/T)_5_ target motifs in the genome or its anticipated function as a transcriptional repressor. Uncertainty also prevails regarding the role of the long known interaction between SALL4 proteins and the Nucleosome Remodelling and Deacetylation (NuRD) complex via a conserved N-terminal peptide ^12,28^. Proteomic analysis showed that SALL4 is present at stoichiometric level comparable to NuRD subunits ^29^, suggesting that NuRD is a major effector of SALL4-mediated repressive activity. Furthermore, NuRD recruitment by SALL4 is reported to be important in the context of iPSC reprogramming ^30^ and cancer cell proliferation ^31^, but the biological relevance of the SALL4-NuRD interaction in stem cells and during development has been questioned ^26,32^.

Here, we have directly addressed these unknowns by high resolution mapping of SALL4 on chromatin and functional characterisation in genetically engineered stem cells and animal models. Our study systematically interrogates SALL4 genome occupancy and its relationship with gene expression. Beyond canonical peak binding we demonstrate more prevalent dispersed binding that reflects base composition. Explainable machine learning predictions and multi-omics investigations established a direct role for SALL4 in modulating chromatin structure via recruitment of the NuRD co-repressor complex. Together, our analyses shed light on an atypical mechanism of gene regulation mediated by SALL4, whereby recognition of frequent AT-rich motifs that are broadly dispersed across transcription units modulate both the epigenome and the transcriptome.

## Results

### SALL4 tracks AT-rich DNA base composition genome-wide

Despite the high frequency of short AT-rich motifs over the genome ^11^, previous reports have only reliably detected SALL4 binding *in vivo* at sites of open chromatin corresponding to enhancer elements ^13,26,27^. As some steps of the ChIP-seq protocol, such as formaldehyde crosslinking and chromatin sonication, may have introduced technical biases that prevent detection of transient interactions ^33–37^, we applied the CUT&RUN protocol ^38^ to map endogenous SALL4 on native chromatin in ESCs. Two independent negative control conditions were adopted to normalise CUT&RUN data: *Sall4* knockout ESCs ^13^ and wildtype ESCs using IgG instead of a specific antibody. The peak detection algorithm MACS2 ^39^ revealed a large number of genomic sites with enriched SALL4 binding (n= 43,973) (Fig. 1A) that overlapped well with published ChIP-seq data ^13^ (Fig. S1A) and carried “active” enhancer marks H3K27ac and H3K4me1 ^40^ (Fig. S1B). Genomic distribution analysis revealed that these sites are predominantly intergenic (48.1%) and intronic (34.8%), with a minority of sites found at promoters (12.7%) (Fig. S1C). Consistent with previous observations ^13^, the base composition of SALL4 peaks is shifted towards AT-rich DNA when compared to CUT&RUN peaks detected in *Sall4* knockout or IgG negative controls (Fig. S1D). However, the SALL4 CUT&RUN data also revealed that the vast majority of mapped reads (91.9%) were localised outside of peaks (Fig. 1B). This provided a first indication that SALL4 might be predominantly dispersed across the genome, rather than being focussed at enhancer elements. Inspection of normalised SALL4 CUT&RUN signal in relation to AT-rich DNA within genomic snapshots (Fig. 1C) strikingly demonstrated that SALL4 binding tracks fluctuations in AT-content over large genomic distances. As a consequence, SALL4 appears to be enriched within large ∼100Kb to megabase-scale blocks of higher AT-content, corresponding to lamina-associated domains (LADs) ^5^ (Fig. 1C, S1E). In contrast, SALL4 is depleted from inter-LADs (iLADs) which are relatively GC-rich. (Fig. 1C, S1E). These observations cannot be explained by a higher frequency of peaks or increased peak signal within AT-rich domains, as artificial exclusion of all peak regions from the normalised SALL4 CUT&RUN signal did not affect the correlation between SALL4 occupancy and AT-rich DNA (Fig. 1C). To further assess this relationship, the entire mouse genome was divided into 10kb windows (Fig. S1F) and sorted according to base composition. SALL4 CUT&RUN signal showed a gradual increase in occupancy proportional to AT-content of the windows (Fig. 1D). Once again, this effect persisted when excluding all peak signal (Fig. S1G). Therefore CUT&RUN analysis uncovered broad genomic binding of SALL4 which is tightly correlated with AT-content. These results reconcile SALL4 chromatin binding *in vivo* with its strong preference for short AT-rich motifs observed *in vitro* ^13,25^ and they raise the possibility that pervasive binding of this protein across the genome (outside of the context of enhancer peaks) has biological relevance.

**Figure 1.**
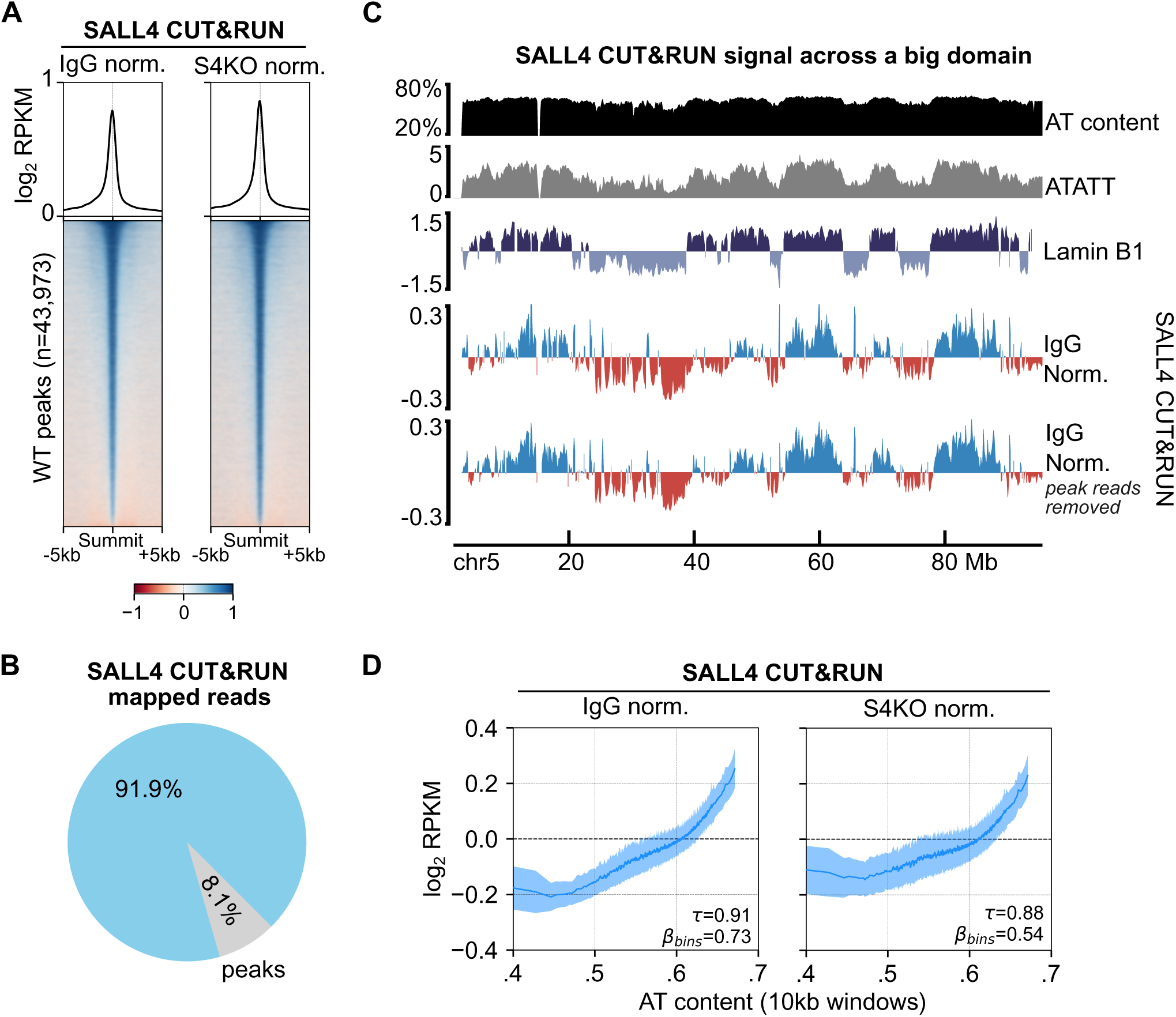
Exploration of the relationship between SALL4 and the AT-rich genome by CUT&RUN analysis. **A.** Heatmap showing normalised SALL4 CUT&RUN signal centred on SALL4 peaks detected in wild-type ESCs (n=43,973). Data was normalised either to SALL4 CUT&RUN in *Sall4* knockout ESCs (S4KO norm.) or using normal IgG in wild-type cells as a negative control (IGG norm.). **B.** Pie chart showing the proportion of SALL4 CUT&RUN reads in wild-type ESCs mapping to the mouse genome, either within (blue) or outside (grey) SALL4 peaks. **C.** Genomic snapshot of mouse chromosome 5 showing global variations in AT-content, short AT-rich motifs (ATATT, preferred SALL4 motif) and SALL4 CUT&RUN signal over large genomic distances. To test the relevance of dispersed SALL4 binding, CUT&RUN reads within peaks were subtracted from total signal. Lamina-associated domains (LADs) and inter-LADs (iLADs) are indicated as genomic features correlating with changes in AT content. **D.** Plots showing normalised SALL4 CUT&RUN across 10kb genomic bins sorted by AT-content. SALL4 signal was smoothed using a rolling mean with a window size of 1000 bins, and values were sampled every 400 bins. Shaded region represent one half standard deviation (SD) error across the 1000 bins.

### Acute degradation of SALL4 correlates gene body occupancy with base composition-dependent transcriptional deregulation

To gain direct mechanistic insight into SALL4 function in ESCs, we generated a degron cell line using the ‘dTAG’ system ^41^ allowing for rapid and controllable depletion of the protein (Fig. 2A). Using CRISPR/Cas9, both *Sall4* alleles were tagged with FKBP12^F36V^ (dTAG), without significantly affecting endogenous protein expression or its localisation at AT-rich pericentric heterochromatin (Fig. S2A, S2B). As expected, tagged SALL4 is rapidly and completely degraded upon acute treatment with dTAG-13 molecule (Fig. 2B). To investigate SALL4-dependent transcriptional regulation, we measured changes in nascent transcription by ‘Transient Transcriptome’ sequencing (TT-seq) ^42,43^ in SALL4-dTAG ESCs treated for 2h with dTAG-13, or with DMSO as a control (Fig. 2A, S2C). Using results from differential expression analysis, and applying a threshold of p-adjusted [padj] value <0.05, we initially divided nascent transcripts into up-regulated (UP) and down-regulated (DOWN) groups compared to a random subset of non-responsive genes expressed in ESCs (padj >0.9 category). UP genes in particular showed higher AT content across their gene units, compared to DOWN genes which were relatively AT-poor (Fig. S2D). Steady-state expression levels differed across categories, DOWN genes were on average more highly expressed than UP genes, with the random set at intermediate levels (Fig. S2D). We next stratified all detectable nascent transcripts into three different subsets depending on thresholds of statistical significance (padj <0.05; 0.05< padj <0.5; 0.5< padj <0.9), using non-responsive genes (padj >0.9) as a negative control panel (Fig. 2C). We reasoned that using multiple significance tiers, rather than a single padj <0.05 cutoff, would potentially capture a continuum of effect sizes reflecting AT-dependent binding of SALL4 to the genome. The magnitude of changes in transcription showed a striking correlation with the AT-content of gene units for all three categories of SALL4-responsive genes (Fig. 2D). This relationship was the strongest for stringently defined SALL4 target genes (padj <0.05), as assessed by Kendall rank correlation (𝜏=0.49) and calculated coefficients from linear regression (βbins=7.64) (Fig. 2D). As our CUT&RUN data showed global binding of SALL4 to AT-rich DNA genome-wide, we decided to investigate the expression of non-coding transcripts such as enhancer RNAs and short intergenic RNAs which are efficiently captured by TT-seq ^42^. To achieve this, transcripts without any known Ensembl annotation ^44^ were analysed (Fig. S2E), revealing that SALL4 depletion also leads to AT-dependent de-regulation of the non-coding transcriptome (Fig. S2F). Strikingly, the great majority of the nascent transcriptome shows base composition dependent transcriptional de-regulation after acute degradation of SALL4.

**Figure 2.**
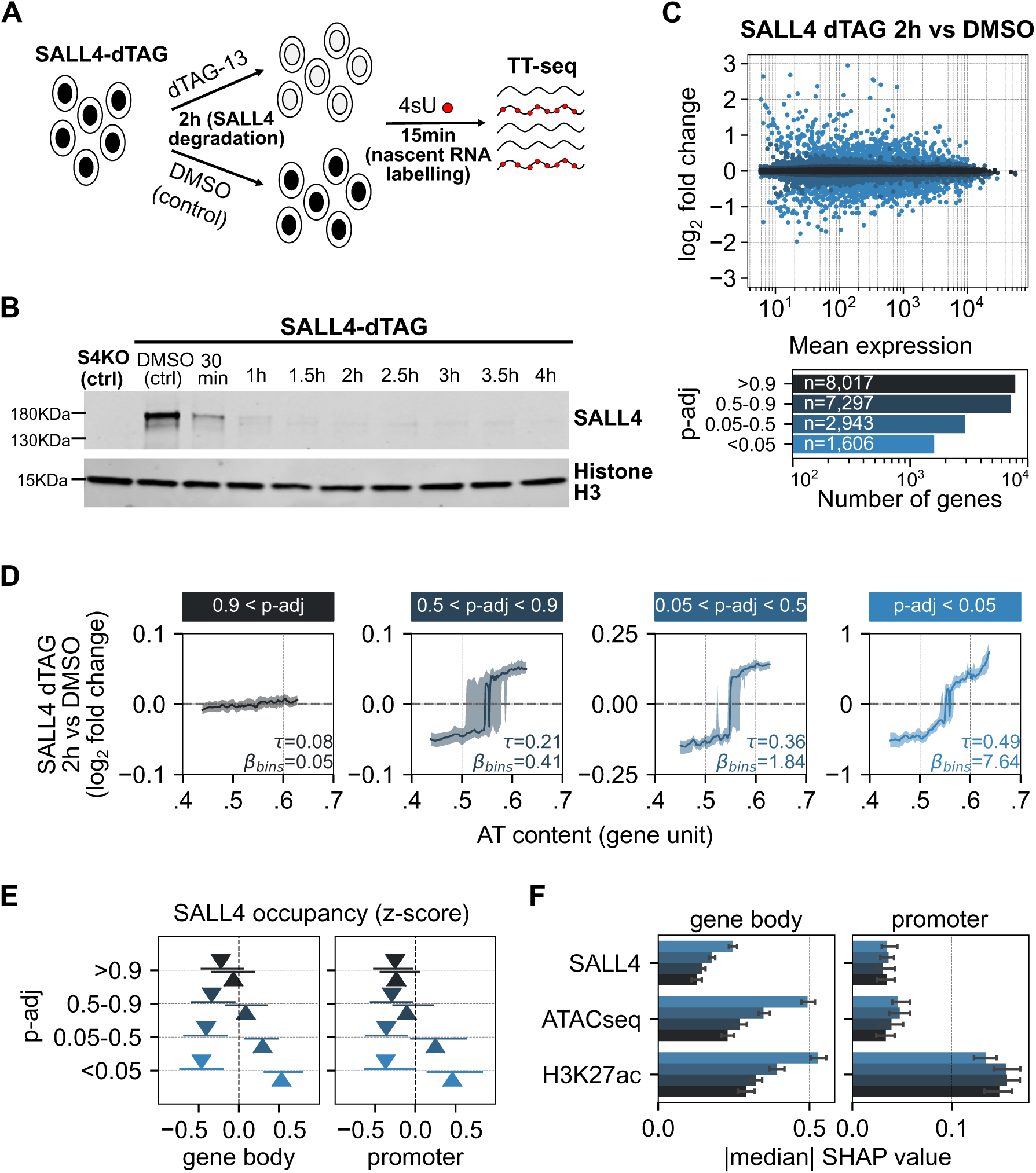
Uncovering direct transcriptional regulation by SALL4 using a rapidly degradable allele. **A.** Diagram summarising the strategy to assess the direct impact of SALL4 on nascent transcription in ESCs, by combining the dTAG degron system with ‘transient transcriptome’ sequencing (TT-seq). **B.** Timecourse Western blot analysis following treatment of SALL4 degron ESCs with dTAG-13. Histone H3 was used as a loading control. Images are representative from two independent replicate experiments. **C.** MA plot showing gene de-regulation following acute degradation of SALL4 in ESCs (2h dTAG-13 versus DMSO). The bar plot shows the number of differentially regulated genes stratified by adjusted p-value (padj) thresholds. **D.** Gene expression changes (log₂ fold change) upon acute SALL4 depletion are plotted similarly across the gene groups in panel C. The solid line represents the rolling median, and the shaded error band represents the inter-percentile range (0.375 to 0.625 quantiles). The strength of association is quantified using Kendall’s correlation coefficient (τ) and the linear regression slope across the bins (β_bins_). **E.** SALL4 CUT&RUN signal (IgG-normalized z-score) at gene bodies (left) and promoters (right) across the gene groups in panel C. Genes are further grouped into up-regulated (up arrow triangle) and down-regulated (down arrow triangle). The plotted triangles correspond to the median SALL4 occupancy and error bars represent the inter-percentile range (0.375 to 0.625 quantiles). **F.** Bar plots showing the absolute median SHAP values from the explainable machine learning model, quantifying the predictive importance of SALL4, ATAC-seq, and H3K27ac features. Importance is shown separately for features derived from gene bodies (left) and promoters (right). Error bars represent the inter-percentile range (0.375 to 0.625 quantiles).

To assess the biological relevance of SALL4 target genes, we performed gene ontology (GO) analysis on the stringent subset of differentially expressed genes identified by TT-seq (padj <0.05). This highlighted multiple developmental pathways and in particular an over-representation of genes associated with neurogenesis among up-regulated genes (Table S1, S2). This observation fits with the hyper-differentiation phenotype observed in SALL4 deficient ESCs ^13,26^. Comparison with previous RNA-seq data from *Sall4* knockout ESCs ^13^ showed an extensive overlap (71%), but target genes identified by TT-seq represent only a small proportion of all differentially expressed genes (≈10%) seen in this mutant cell line (Fig. S2G). It is likely that additional transcriptional defects seen in cells that have adapted to the absence of SALL4 over many generations mainly correspond to secondary and/or compensatory effects ^45–47^. Therefore, our experimental approach combining a SALL4 degron allele with nascent transcriptomics allowed for the identification of likely primary target genes, which can be directly linked with phenotypic consequences associated with loss of this protein.

### Explainable machine learning predicts the transcriptional impact of SALL4 and chromatin structure over target gene bodies

Having established widespread base composition dependent transcriptional regulation after acute SALL4 loss (Fig. 2D), we predicted that this transcription factor would cover entire gene units via pervasive binding to AT-rich DNA (Fig. 1D). To address this hypothesis, we quantified SALL4 CUT&RUN signal over gene units, dividing promoter and gene body regions (Fig. S2H). This data was visualised for the different categories of target genes identified by TT-seq (see Fig. 2C), showing that SALL4 CUT&RUN signal was consistently above average (z-score >0) for up-regulated target genes in categories with padj <0.9, and below average (z-score <0) for all down-regulated target genes (Fig. 2E) at both gene bodies and promoters. These trends persisted after artificially excluding (intragenic) SALL4 peaks from analysis (Fig. S2I), suggesting that dispersed binding of this transcription factor is primarily responsible for these relationships. It is notable that TT-seq analysis in SALL4 degron ESCs highlighted similar numbers of genes being up- and down-regulated by this transcription factor (Fig. 2C). Our data is compatible with a role for SALL4 as a repressor, as SALL4 chromatin binding over gene units directly correlates with transcriptional up-regulation after acute SALL4 depletion (Fig. 2E, S2I). In contrast, SALL4 CUT&RUN signal is paradoxically depleted over down-regulated targets, implying that these genes may not be directly activated by SALL4 (Fig. 2E, S2I).

To independently assess the importance of SALL4 binding and other chromatin features across gene bodies and promoters, we utilised an explainable machine learning framework ^48^. Using an accurate deep learning model trained on a combination of SALL4 CUT&RUN and multiple other epigenomic datasets as inputs (Table S3) ^48^, we predicted RNA PolII occupancy in wild-type ESCs using per-feature total signal over promoters and gene bodies (Fig. S2J). We applied Shapley Additive exPlanations (SHAP) to disentangle the respective contributions of each feature to ongoing transcription (RNA PolII occupancy over individual genes), and stratified genes according to TT-seq results in SALL4 degron ESCs as described above (see Fig. 2C). Our analysis revealed that, in addition to SALL4 occupancy, both chromatin accessibility (ATAC-seq) and histone acetylation (H3K27ac ChIP-seq) signal across gene bodies were consistently influential predictors of transcription in wild-type ESCs, and their relative importance tracked the sensitivity of genes to SALL4 degradation (Fig. 2F). In contrast, when promoter regions were analysed, H3K27ac signal appeared predominant and no feature showed preferential importance at SALL4-responsive genes compared to non-responsive genes (Fig. 2F). Together, these predictions support the hypothesis that chromatin structure over gene bodies, rather than promoters, is a key determinant of SALL4-dependent transcriptional regulation in ESCs.

### SALL4 depletion rapidly alters chromatin structure independent of transcription

Modelling suggests an interdependency between SALL4 occupancy and chromatin structure in stem cells. To test this experimentally, we performed a timecourse of ATAC-seq and H3K27ac CUT&RUN in SALL4-dTAG ESCs at 0.5h, 3h and 9h after acute SALL4 depletion (Fig. 3A, S3A). Comparing dTAG and DMSO (control) treated samples using standard peak-calling algorithms (Fig. S3B) showed that acute degradation of SALL4 causes widespread gains in chromatin accessibility within 0.5h (∼21,000 out of a total of ∼162,000 ATAC-seq peaks). Similar results were obtained at 3h and 9h post-degradation (∼28,000 peaks and ∼26,000 peaks, respectively; Fig. S3B). In contrast, H3K27ac-positive peaks showed negligible changes across the entire timecourse (0, 39 and 229 changed loci at 0.5h, 3h and 9h, respectively) indicating that enhancer/promoter acetylation levels remained largely unchanged after degradation of SALL4 (Fig. S3B). As expected, SALL4 peaks detected by CUT&RUN (Fig 1A) correspond to a subset of all H3K27ac and ATAC-seq peaks in ESCs (Fig. S3C) ^13^, but unexpectedly the vast majority of differentially accessible peaks after acute SALL4 depletion (0.5h) neither coincided with SALL4 CUT&RUN peaks (see Fig. S3D) nor overlapped convincingly with direct target genes identified by TT-seq (Fig S3D). Therefore, the functional significance of increased accessibility within peaks in response to SALL4 degradation is unclear. On the other hand, we noticed that a large fraction of ATAC-seq and H3K27ac CUT&RUN reads were located outside regions designated as peaks (41% and 89.6%, respectively; Fig. S3E). In line with predictions from our deep learning model, this raised the possibility that epigenomic signal broadly distributed along gene bodies may have more functional relevance (Fig. 2F).

**Figure 3.**
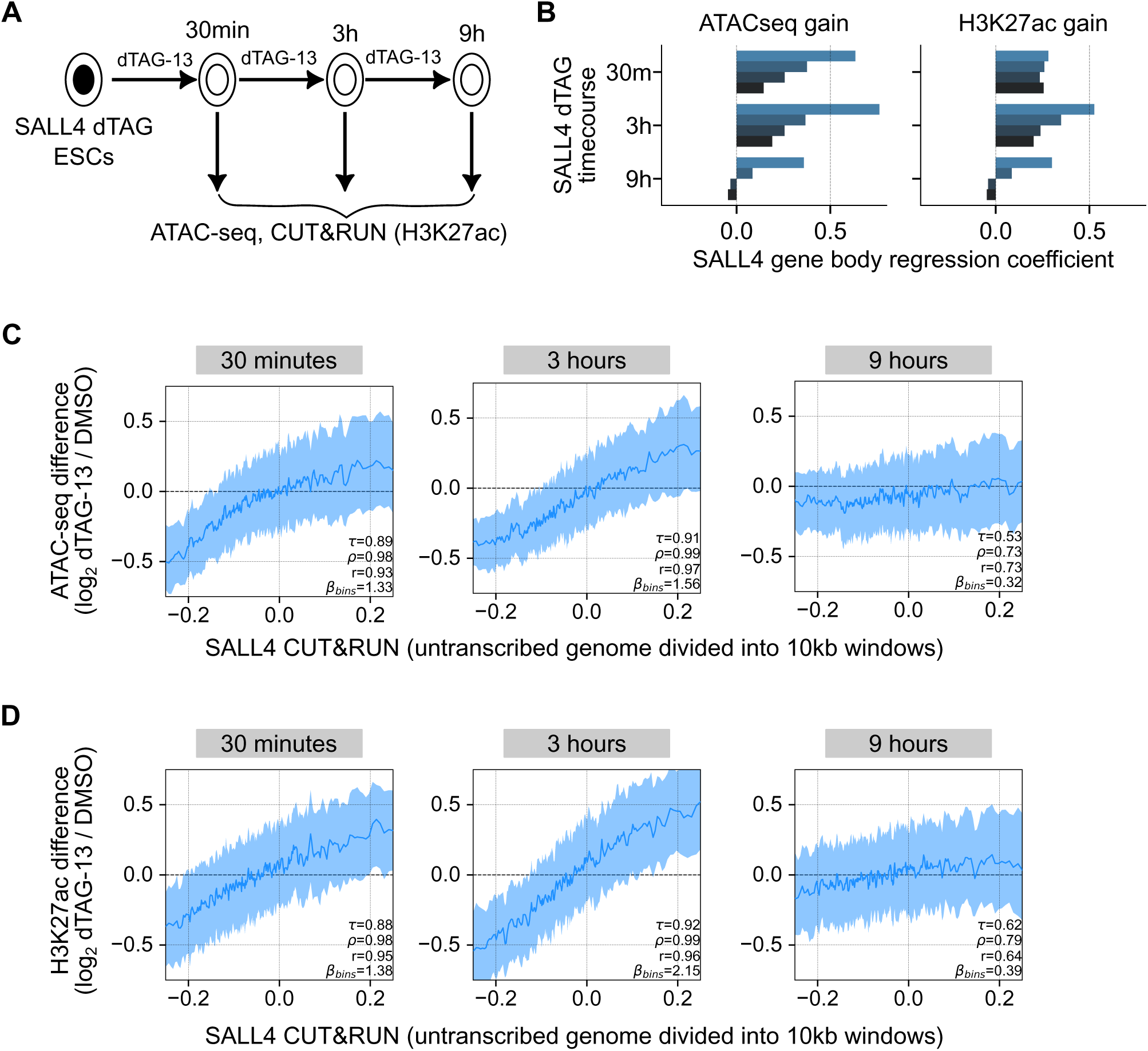
Investigation of chromatin changes directly mediated by SALL4. **A.** Diagram summarising the experimental strategy for the SALL4-dTAG timecourse. SALL4-dTAG ESCs were treated with dTAG-13 for 30 minutes, 3 hours, and 9 hours, followed by ATAC-seq and H3K27ac CUT&RUN analysis. **B.** Bar plots of linear regression slopes across the bins (β_bins_) when gene body gain of ATAC-seq (left) or H3K27ac (right) is regressed against SALL4 occupancy. The colours represent the set of differentially regulated SALL4 target gene categories in Fig. 2C. Rolling median plots visualising changes in **C.** ATAC-seq and **D.** H3K27ac at transcriptionally inactive 10kb windows at 0.5h, 3h and 9h post SALL4 degradation with respect to SALL4 CUT&RUN occupancy in WT mESCs. The shaded band represents the inter-percentile range (0.375 to 0.625 quantiles). Correlation (Kendall’s τ, Pearson’s r) and linear regression slope (β_bins_) are shown for each plot.

To test whether dispersed ATAC-seq and H3K27ac signal is informative in interpreting the consequences of SALL4 degradation on gene expression, we used simple linear regression to quantify the relationship between SALL4 CUT&RUN signal (in unperturbed wild-type ESCs) and changes in chromatin accessibility or histone acetylation over entire gene units (Fig. S3F, S3G). Focussing on gene body signal, we observed a predominant gain in ATAC-seq signal over SALL4 transcriptional targets, with regression coefficients mirroring their sensitivity to SALL4 degradation (Fig. 3B). The magnitude of the changes was maximal 0.5 to 3h after SALL4 degradation but somewhat reduced after 9h, suggesting relatively rapid compensatory mechanisms at the level of chromatin (Fig. 3B). Similar results were obtained using H3K27ac CUT&RUN data, although the relationship between gain in histone acetylation and SALL4 occupancy over target gene bodies only became evident after 3h (Fig. 3B). Unlike gene bodies, promoter regions showed small regression coefficients in relation to ATAC-seq and H3K27ac signal and no consistent relationship with SALL4 target genes (Fig. S3H). These experimental findings are consistent with model predictions generated using epigenomic data in unperturbed wild-type ESCs (Fig. 2F). In particular, regression coefficients calculated over gene bodies from ATAC-seq/H3K27ac CUT&RUN data in SALL4 degron cells mirror SHAP values extrapolated from our deep learning model (Fig. 2F, 3B). Overall, the combined investigations from modelling and experiments indicate a clear relationship between SALL4 activity and chromatin structure over transcription units.

While these results raise the possibility that SALL4 directly promotes chromatin compaction and histone de-acetylation, it is well-known that transcription activity itself influences the epigenome ^49–52^ making assignment of causality uncertain. To address this issue, we took advantage of our nascent transcriptomics data (TT-seq) which detected all transcription units in ESCs (Fig. S2E) and thereby defined a complementary set of transcriptionally inactive regions. We defined these regions as those lacking detectable nascent transcription in both control (DMSO) and SALL4-depleted (dTAG-13) conditions, and which also fell outside of all annotated transcription units. The transcriptionally inactive genome was divided into 10kb windows sorted based on levels of SALL4 CUT&RUN signal, allowing us to test whether loss of SALL4 might induce chromatin changes in the absence of transcription. This showed a striking relationship between SALL4 occupancy and increased chromatin accessibility (ATAC-seq) and histone acetylation (H3K27ac CUT&RUN) following SALL4 depletion (Fig. 3C, 3D). As observed with transcribed genes, these effects were significantly attenuated 9h post-degradation indicating compensatory effects at this later time point (Fig. 3C, 3D). Our results at transcriptionally silent loci closely match epigenomic changes observed over the bodies of SALL4 target genes (Fig. S3I, S3J). We conclude that the immediate gains in genomic accessibility and histone acetylation observed after degradation of SALL4 are not a secondary consequence of changed transcriptional activity, but result from a direct influence of the SALL4 transcription factor upon chromatin structure.

### Recruitment of the NuRD co-repressor complex is essential for SALL4 function in pluripotent cells and during development

The experimental and computational approaches above suggest that histone de-acetylation and chromatin inaccessibility are key molecular consequences of SALL4 binding to DNA. A potential link between SALL4 and these features of the epigenome is the NuRD co-repressor complex, which interacts with SALL4 via an evolutionarily conserved N-terminal peptide (NT) recognising the WD40 domain of RBBP4/7 ^28,53^ (Fig. 4A, S4A). NuRD includes both histone deacetylase and nucleosome remodelling activities and its lack of recruitment could therefore explain the short-term epigenomic consequences of sudden SALL4 degradation (see Fig. 3). To directly test this hypothesis, we used CRISPR/Cas9 in ESCs to generate two mutant forms of SALL4 that specifically disrupt NuRD interaction (Fig. 4A): “NTΔ” in which core amino acids within the NT domain were deleted; and “NTmut” in which the key RRK motif ^31^ was substituted by three alanines. As a control, we generated a “recovery” CRISPR line containing silent mutations in several codons within the NT (WTrec) and therefore expressing a wild-type protein. First, we confirmed stable SALL4 expression by Western blot analysis, which established near wild-type levels in all cell lines (Fig. S4B). As expected, co-immunoprecipitation confirmed a significant loss of interaction between SALL4 and the NuRD subunit MTA1 in both NTmut and NTΔ ESCs (Fig. S4C). SALL4 immunoprecipitation combined with label-free mass spectrometry analysis (IP/MS) (Fig. S4D) demonstrated a >4-fold depletion of all core NuRD subunits, as well as NuRD-associated proteins CDK2AP1 ^54^ and BEND3 ^55^, (Fig. 4B). In contrast, hetero-dimerisation with other SALL family proteins SALL1/2/3 ^56,57^ was largely unaffected by mutation of the NT domain (Fig. 4B). As SALL4 is reportedly highly enriched within NuRD in ESCs ^29^, we asked whether the overall stability or composition of this co-repressor complex was affected in our mutant cell line. IP/MS of the core NuRD subunit MBD3 ^58,59^ (Fig. S4E) did not reveal significant changes in NuRD composition in NTmut ESCs, apart from the expected loss of SALL4 (Fig. S4F). Therefore, mutation of the NT domain specifically disrupts the interaction between SALL4 and NuRD, without affecting the overall structure of the co-repressor or its SALL4-independent functions^60,61^.

**Figure 4.**
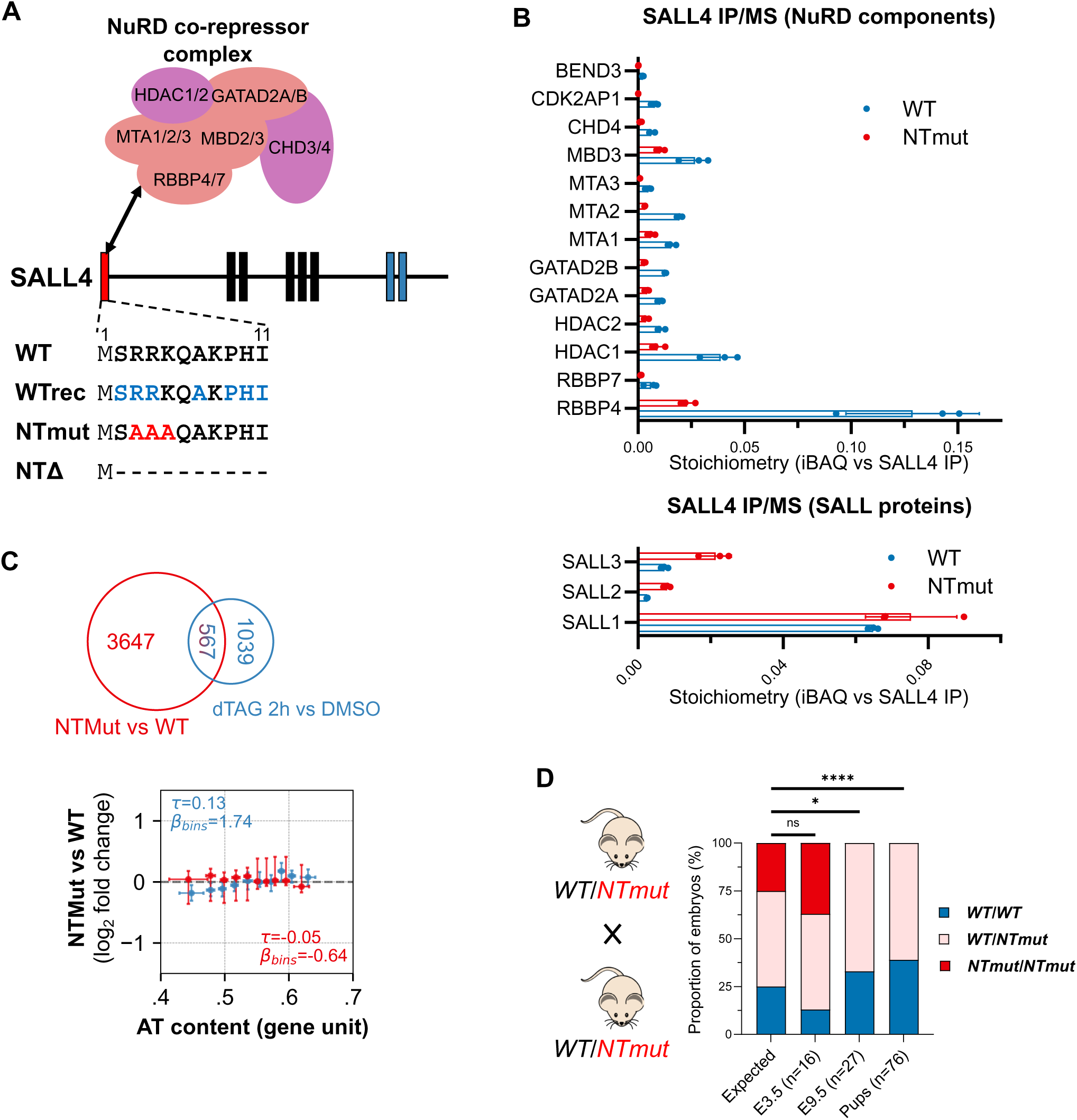
Molecular and phenotypic consequences of disrupting the interaction between SALL4 and NuRD co-repressor. **A.** Diagram showing mutations introduced by CRISPR/Cas9 at the N-terminus of SALL4 coding sequence. To disrupt the interaction with NuRD co-repressor complex, key residues interacting with RBBP4/7 were mutated to alanine (NTmut) or the entire interaction domain was deleted (NTΔ). The “recovery” allele (WTrec) contains silent mutations in highlighted residues, thereby generating a wild-type protein sequence. **B.** Relative stoichiometries of NuRD subunits and SALL proteins following SALL4 IP/MS in wild-type or NTmut ESCs. Absolute protein amounts were estimated using iBAQ intensities and normalised to SALL4 IP in each sample. Each data point represents an independent replicate experiment (n=3). Error bars: Standard deviation (SD). **C.** Venn diagram showing the overlap of differentially expressed genes (compared to wild-type) in NTmut and NTΔ ESCs, as assessed by RNA-seq analysis with direct targets of SALL4, as assessed by TT-seq analysis. Gene expression changes (log₂ fold change) in NTmut/Δ ESCs across genes grouped in increasing bins of AT content and error bars represent the inter percentile ranges (0.375 to 0.625 quantiles). For each gene set, the strength of association is quantified using Kendall’s 𝜏 and the linear regression slope across the bins (β_bins_). **D.** Bar graphs showing the genotype of embryos (E3.5, E9.5) and live pups (P21) following crosses between heterozygous *WT/NTmut* mice. Statistical comparisons with expected Mendelian ratios were performed using the Fisher’s exact test. Homozygous mutation of the NuRD interaction domain is completely lethal by embryonic day E9.5.

To assess the phenotypic consequences of disrupting the SALL4-NuRD interaction, we performed RNA-seq analysis in all our mutant cell lines (WTrec ESCs were considered as independent wild-type clones to account for clonal variability). Mutation of the NT domain of SALL4 (NTmut/Δ) resulted in several thousand differentially expressed genes indicating a global change in the transcriptome (Fig. 4C). Changed transcripts showed a poor overlap with primary target genes of SALL4 identified by TT-seq in degron ESCs, and their level of de-regulation showed no relationship with AT-content (Fig. 4C). These observations raised the possibility that the transcriptional defects observed in NTmut/Δ cell lines reflect an altered cell state rather than a direct consequence of SALL4 loss-of-function. Indeed, gene ontology analysis on differentially expressed genes highlighted multiple developmental pathways associated with organogenesis, suggesting compromised pluripotency (Table S4, S5). To test whether NuRD recruitment is required for embryonic development *in vivo*, we generated transgenic mice with mutation in the SALL4 NT domain (NTmut) by CRISPR/Cas9 injection of fertilised zygotes ^62^. Following crosses of heterozygous *NTmut* animals, some embryos were collected at embryonic days E3.5 and E9.5, and living pups were genotyped 21 days after birth. These experiments established that homozygous mutation of the NT domain was embryonically lethal shortly after implantation by E9.5 (Fig. 4D, Table 1), thus mimicking the phenotype observed after a complete *Sall4* knockout ^14,15^. Together, these findings demonstrate that recruitment of NuRD co-repressor complex is essential for SALL4 biological function in stem cells and during early embryonic development.

**Table 1.**
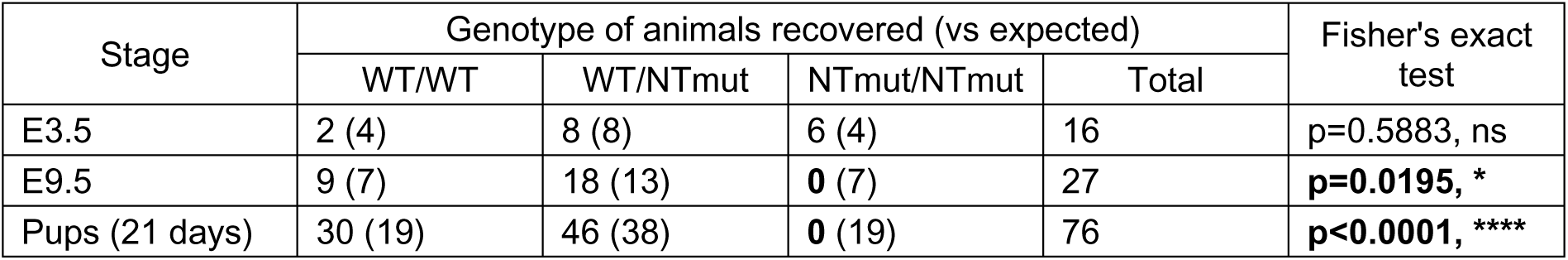
Genotypic analysis of progeny from NTmut heterozygous crosses.

| Stage | Genotype of animals recovered (vs expected) |  |  |  | Fisher's exact test |
| --- | --- | --- | --- | --- | --- |
|  | WT/WT | WT/NTmut | NTmut/NTmut | Total |  |
| E3.5 | 2 (4) | 8 (8) | 6 (4) | 16 | p=0.5883, ns |
| E9.5 | 9 (7) | 18 (13) | **0** (7) | 27 | **p=0.0195, \*** |
| Pups (21 days) | 30 (19) | 46 (38) | **0** (19) | 76 | **p<0.0001, \*\*\*\*** |

### Artificial recruitment of a different co-repressor indicates that chromatin remodelling activity is essential for SALL4 function

Multiple co-repressor complexes are expressed in ESCs, and often share the same subunits with scaffolding or enzymatic activities ^60,63^. Among them, Sin3 co-repressor includes RBBP4/7 and histone deacetylases HDAC1/2 found in NuRD but lacks nucleosome remodelling subunits CHD3/4 (Fig. 5A). To test whether a different repressive complex without chromatin remodelling activity can sustain SALL4 biological function, we designed a domain swap aimed at removing the NuRD-interaction domain of endogenous SALL4 and replacing it with the evolutionarily conserved Sin3-interaction domain (SID) of MAD1/MXD1 protein ^64–70^ (Fig. 5A and S5A). Western blot analysis confirmed that homozygous SID ESCs expressed SALL4 close to levels observed in wild-type cells (Fig. S5B). As expected, immunoprecipitation of SALL4 SID followed by either Western blot (Fig. S5C) or mass spectrometry analysis (Fig. 5B, S5D) showed a dramatic loss of interaction with NuRD-specific subunits (similar to NTmut), accompanied by new protein-protein interactions with all known Sin3 subunits and associated proteins in ESC (SIN3A, SAP30/L, SAP130, FAM60A, SUDS3,BRMS1/L, ING1/2, ARID4A/B, TET1) ^71^. Proteins found in both NuRD and Sin3 complexes (RBBP4/7, HDAC1/2) showed similar interaction between SALL4 SID and wild-type proteins, and heterodimerisation with other SALL proteins appeared unaffected (Fig. S5E). To assess the biochemical activity of co-repressors recruited by SALL4 in wild-type and mutant conditions, we used a chemiluminescent reporter assay for HDAC1/2 function in SALL4 immunoprecipitates. While mutation of the NuRD-interaction domain (NTmut/Δ ESCs) caused a significant decrease in histone deacetylase activity, SALL4 SID recovered enzymatic activity near wild-type levels (Fig. 5C). Together, these analyses demonstrate that a domain swap of SALL4 N-terminal domain with a SID peptide is sufficient to prevent its interaction with the NuRD co-repressor and instead cause recruitment of the Sin3 corepressor complex.

**Figure 5.**
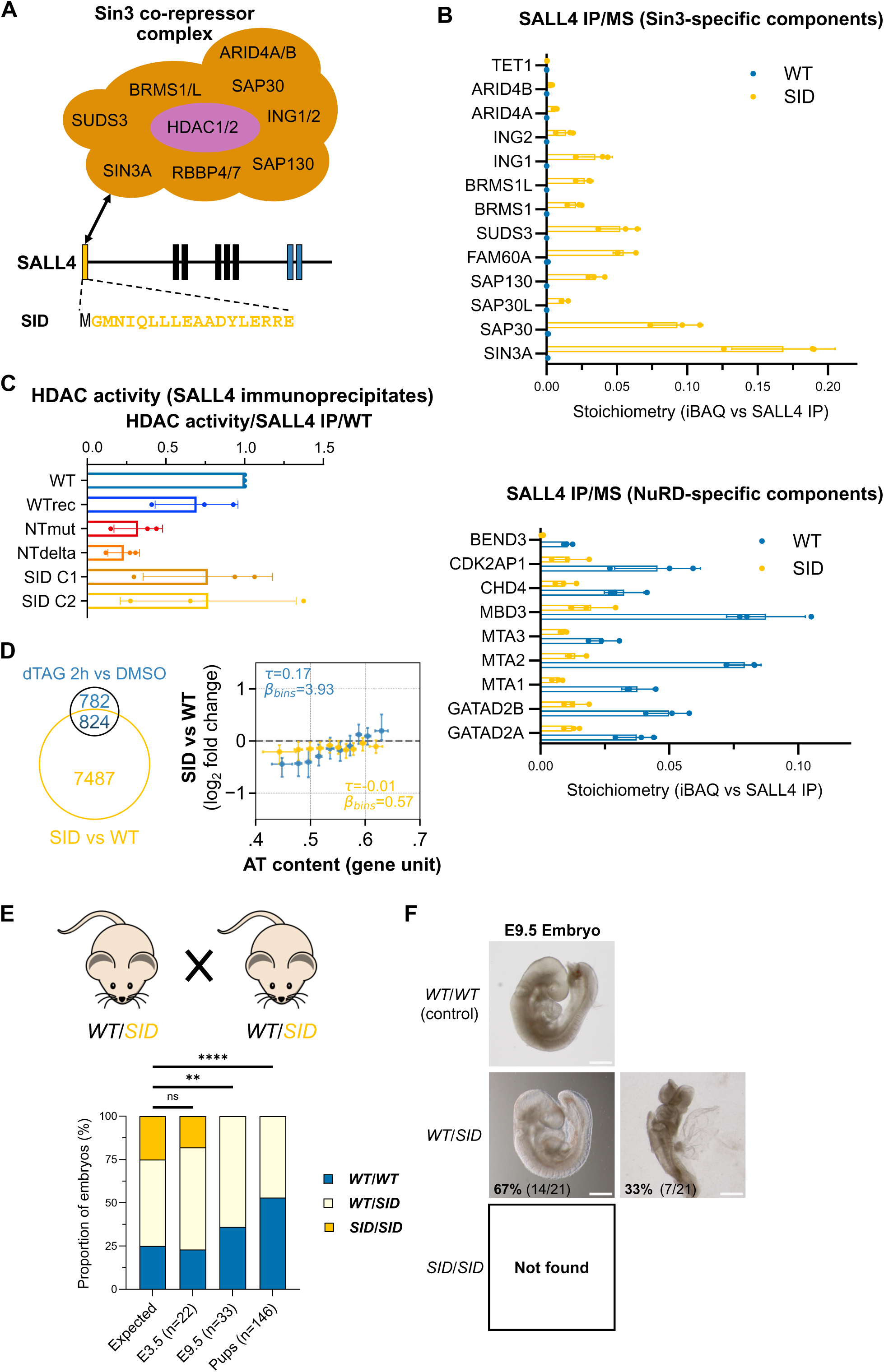
Domain swap experiment to artificially substitute NuRD with Sin3 co-repressor at SALL4 targets. **A.** Diagram showing mutations introduced by CRISPR/Cas9 at the N-terminus of SALL4 coding sequence. The NuRD interaction domain of SALL4 (residues 2-11) was deleted and replaced with the Sin3-interaction domain of the transcriptional repressor MAD1 (residues 6-23). This domain swap is expected to result in artificial recruitment of Sin3 co-repressor by endogenous SALL4. **B.** Relative stoichiometries of NuRD-specific, Sin3-specific subunits following SALL4 IP/MS in wild-type or SID ESCs. Absolute protein amounts were estimated using iBAQ intensities and normalised to SALL4 IP in each sample. Each data point represents an independent replicate experiment (n=3). Error bars: Standard deviation (SD). **C.** Bar graph showing the histone deacetylase activity measured in SALL4 immunoprecipitates from wild-type and mutant ESC lines. Measurements were normalised to SALL4 protein amounts (determined by Western blot) and expressed relative to wild-type. Each data point represents an independent replicate experiment (n=3). Error bars: Standard deviation (SD). **D. C.** Venn diagram showing the overlap of differentially expressed genes (compared to wild-type) in SID ESCs, as assessed by RNA-seq analysis with direct targets of SALL4, as assessed by TT-seq analysis. Gene expression changes (log₂ fold change) in SID ESCs across genes grouped in increasing bins of AT content and error bars represent the inter percentile ranges (0.375 to 0.625 quantiles). For each gene set, the strength of association is quantified using Kendall’s 𝜏 and the linear regression slope across the bins (β_bins_). **E.** Bar graphs showing the genotype of embryos (E3.5, E9.5) and live pups (P21) following crosses between heterozygous *WT/SID* mice. Statistical comparisons with expected Mendelian ratios were performed using the Fisher’s exact test. Homozygous SID domain swap is completely lethal by embryonic day E9.5. **F.** Representative images of E9.5 embryos from indicated genotypes, obtained by crossing heterozygous *WT/SID* mice. Heterozygous SID embryos presented either a normal morphology, or severe morphological abnormalities (including defects in neural tube closure). Proportion of embryos of a given phenotype is indicated directly on the picture.

To evaluate the phenotypic consequences of this domain swap, we performed RNA-seq in several SALL4 SID ESC clones. This analysis revealed a profound de-regulation of the transcriptome compared to wild-type ESCs (>8,000 de-regulated transcripts). Once again, differentially expressed genes showed a poor overlap with direct target genes of SALL4 and a very weak correlation with AT-content (Fig. 5D), indicating the predominance of compensatory or adaptative mechanisms. Nevertheless, comparison with RNA-seq data from NTmut/Δ ESCs indicated a partial recovery of transcriptional defects caused by loss of NuRD recruitment (1,716 genes), but also new gene de-regulation resulting from artificial recruitment of Sin3 complex (5,813 genes) (Fig. S5F). GO term analysis identified de-regulation of genes associated with developmental pathways but also with general chromatin and transcriptional processes, indicating an altered stem cell state (Table S6, S7). To determine whether developmental defects observed after loss of NuRD interaction could be partially reverted by Sin3 recruitment, we generated a SALL4 SID transgenic mouse model. Similar to results obtained in the NTmut line, homozygous SID mutation had a fully penetrant lethal phenotype as early as embryonic day E9.5 (Fig. 5E, Table 2). Furthermore, we noticed a bias towards a wild-type genotype in live pups compared to expected Mendelian ratios (Fig. 5E, S5G, Table 3), indicating that heterozygous SID mutation exerts a dominant-negative effect leading to partially penetrant embryonic lethality. Accordingly, we observed gross morphological abnormalities (mainly failure of neural tube closure) in ≈33% of heterozygous SID embryos collected at E9.5 (Fig. 5F). Our results demonstrate that the NuRD complex recruitment is indispensable for SALL4 biological function and cannot be substituted by Sin3.

**Table 2.** Genotypic analysis of progeny from SID heterozygous crosses.

| Stage | Genotype of animals recovered (vs expected) |  |  |  | Fisher's exact test |
| --- | --- | --- | --- | --- | --- |
|  | WT/WT | WT/SID | SID/SID | Total |  |
| E3.5 | 5 (5) | 13 (12) | 4 (5) | 22 | p>0.9999, ns |
| E9.5 | 12 (8) | 21 (17) | **0** (8) | 33 | **p=0.0090, \*\*** |
| Pups (21 days) | 77 (36) | 69 (74) | **0** (36) | 146 | **p<0.0001, \*\*\*\*** |

**Table 3.** Genotypic analysis of progeny from WT and SID heterozygous crosses.

| Stage | Genotype of animals recovered (vs expected) |  |  | Fisher's exact test |
| --- | --- | --- | --- | --- |
|  | WT/WT | WT/SID | Total |  |
| Pups (21 days) | 163 (103) | 42 (103) | 205 | **p<0.0001, \*\*\*\*** |

## Discussion

Transcription factors are traditionally assumed to regulate gene expression by acting at specific genomic locations, corresponding to regulatory elements ^72^. However, enrichment of transcription factors at enhancers and promoters is usually poorly predictive of transcriptional output ^73–76^, highlighting our incomplete understanding of this fundamental biological process. Expanding this view of gene regulation, we found that SALL4 acts as a “global” modulator of gene expression via promiscuous binding to short A/T motifs whose frequency reflects local DNA base composition. How might dispersed binding of SALL4 modulate gene expression? Previous *in vitro* studies suggested that the A/T motifs bound by the DNA-binding domain of SALL4 (ZFC4) are ∼5 nucleotides in length ^13,20–22^, which means that on the bulk genome (∼60% A/T) about 8x binding sites would be expected every 100 base pairs. Therefore, an average gene of 30kb with a typical 60% A/T content would possess over 2,300 potential target sites, whereas genes of the same length that were 50% or 40% A/T would have 937 and 307 target motifs, respectively. We hypothesise that these differences result in variable densities of bound SALL4, which in turn recruit variable amounts of NuRD leading to chromatin compaction and histone deacetylation in proportion to their A/T content. Extended “blanket” repression of this kind may influence gene expression by affecting RNA polymerase elongation or some other aspect of the transcription process.

In order to determine the context-dependent importance of SALL4 and NuRD-associated chromatin features in influencing transcription, we utilised explainable deep learning models trained on data profiled from wild-type ESCs. In addition to SALL4 genomic occupancy, ATAC-seq and H3K27ac signal across gene bodies appeared as important determinants in predicting the expression of SALL4 target genes. Expanding on these predictions, multi-omic analyses in SALL4 degron cells revealed that this transcription factor has a direct impact on the epigenome. Interestingly, degradation of SALL4 results in a rapid increase in histone acetylation and chromatin accessibility over entire gene units corresponding to SALL4 transcriptional targets. These changes were not only proportional to SALL4 CUT&RUN signal, but also to the degree of transcriptional upregulation following degradation of the protein. Importantly, this effect on chromatin structure was not confined to transcription units but also applied to intergenic domains where no nascent transcripts were detected. Our finding that such inert regions of the genome responded in the same way as genes argues strongly that these effects are not a secondary consequence of altered gene expression, which is notoriously inter-connected with chromatin accessibility and histone acetylation status ^49–52^.

Prior emphasis on SALL4’s potential role at enhancers is explained by evidence from ChIP-seq experiments that SALL4 primarily localises at active enhancers in ESCs ^13,26,27^. In contrast, our CUT&RUN data shows predominantly dispersed binding across the genome (>90% of mapped reads). SALL4 enrichment is proportional to AT-content, presumably due to recognition of short AT-rich motifs via zinc finger cluster ZFC4, which we have shown previously to be essential for SALL4 function ^13^. We suggest that detection of transient interactions with AT-rich DNA may have been prevented in past studies by technical limitations of ChIP-sequencing ^33–37^. Furthermore, as chromatin analyses usually rely on peak-calling algorithms, SALL4 reads outside of peak regions may have been dismissed as “background” signal. While our analyses support a key role for SALL4 over gene bodies, we cannot exclude that localisation of this transcription factor elsewhere can also have a significant impact on transcription. In common with other studies on a variety of transcription factors ^73–76^, our attempts to link SALL4 CUT&RUN peaks with transcriptional de-regulation occurring in SALL4 degron cells were unsuccessful. At face value, this implies that SALL4 binding at these sites of open chromatin are of minimal functional relevance. It remains possible, however, that aspects of our analysis are sub-optimal. For example, linking enhancers with their candidate target genes based on genomic distance may be too simplistic, as enhancers are known to mediate long-range chromatin looping ^77^. Interestingly, at a large genomic scale SALL4 is enriched over lamina-associated domains, which are preferentially located close to the nuclear membrane and correspond to repressive chromatin compartments characterised by high AT-content ^5^. Whether SALL4 influences sub-nuclear genome architecture has not so far been investigated.

In addition to transcriptional up-regulation, many genes produced fewer transcripts after SALL4 degradation. Superficially, this implies that their native expression levels are somehow sustained by this protein. However, both SALL4 occupancy and AT-content are anticorrelated with down-regulation, arguing against the notion that SALL4 can act as a transcriptional activator. A plausible alternative hypothesis is that SALL4 loss leads to rapid redistribution of the transcriptional machinery to AT-rich genes at the expense of GC-rich genes which have on average high basal expression levels ^5,78^. A precedent for such disruption of global chromatin equilibrium has been reported in the case of SWI/SNF which appears to indirectly promote Polycomb repression by preventing its pervasive re-distribution to non-target genomic loci ^79^.

Considering its well characterised biochemical interaction with SALL4 ^28,53^, the NuRD co-repressor is a prime candidate to mediate this function via histone deacetylation due to HDAC1/2 and nucleosome remodelling due to CHD3/4 ^60,80^. To test the functional relevance of NuRD recruitment, we introduced inactivating mutations within the N-terminal domain SALL4 ^31^. This allowed us to specifically disrupt protein-protein interactions with NuRD subunits without affecting other functional domains within the SALL4 protein. Homozygous mutation of the NuRD-interaction domain caused alterations in the stem cell state as well as peri-implantation lethality in mouse embryos, similar to a complete *Sall4* knockout ^14,15^. We artificially swapped SALL4 N-terminal domain with a Sin3-interaction domain (SID) in order to recruit a co-repressor with the same histone deacetylase composition but lacking chromatin remodelling activity ^70^. This mutation resulted in major transcriptional defects in ESCs and was incompatible with embryonic development in mice. These findings highlight the fact that NuRD recruitment is indispensable for SALL4 function. The reason why Sin3 fails to complement is unknown, but one possibility is that the nucleosome remodelling capability normally mediated by CHD3/4 is essential for regulation via SALL4. Notably, the N-terminal NuRD interaction domain is present in all vertebrate SALL proteins ^12^, suggesting that recruitment of this corepressor is an essential and evolutionarily conserved function of all these proteins. In line with this view, a recent study identified developmental defects in mice carrying mutations in the NuRD-interaction domain of SALL1 ^81^.

In summary, we show that dispersed binding of SALL4 to AT-rich DNA and subsequent recruitment of NuRD co-repressor mediates “fine-tuning” of gene expression in ESCs. Other factors are known to bind promiscuously to AT-rich DNA, including AT-Hook proteins HMGA1/2 ^82^ and AT-rich interacting domain proteins (ARIDs) ^83^. However, due to their relatively relaxed sequence specificity, these factors are usually considered to only play a supportive role in transcriptional regulation. Based on our progress in understanding SALL4 function, it may be of interest to re-visit the role of other AT-binding proteins as potential regulators of the global transcriptome and epigenome.

## Materials and methods

### Experimental procedures

#### Animal work (mice)

The *Sall4 NTmut* mouse line was generated by injection of ribonucleoprotein complexes (RNPs) into WT C57BL/6J zygotes for CRISPR/Cas9 genome editing. RNPs were dissolved in 0.1x TE buffer and contained Cas9 protein, a synthetic guide RNA (sgRNA) targeting *Sall4* Start codon and a single-stranded DNA template for homologous recombination carrying mutation in the NuRD-interaction domain (reagents ordered from Integrated DNA Technologies). Genotyping was performed in the resulting offspring by PCR amplification of *Sall4* alleles (see primers) and Sanger sequencing. Male and female animals with the desired mutation (NTmut) were crossed with WT C57BL/6J animals at 7-8 weeks. Animals were routinely genotyped by PCR combined with restriction fragment length polymorphism (RFLP) analysis, using restriction enzymes cutting specifically the wild-type (PspFI ThermoFisher Scientific cat. FD2224) or NTmut allele (AlwNI, New England Biolabs cat. R0514S), respectively. Heterozygotes identified from these crosses were inter-crossed to generate homozygotes. Heterozygotes were also crossed with WT C57BL/6J to maintain the line.

The *Sall4 SID* mouse line was generated by injection of heterozygous SID ESCs (see section below for genetic engineering in cell lines) into WT C57BL/6J mouse blastocysts using standard methods. Resultant male and female chimeras were crossed with WT C57BL/6J animals at 7-8 weeks and coat colour was used to identify germline offspring. Transmission of the targeted *Sall4* allele was confirmed by PCR (see primers) and Sanger sequencing. Heterozygotes identified from these crosses were inter-crossed to generate homozygotes. Animals were routinely genotyped by PCR combined with RFLP analysis, using restriction enzymes cutting specifically the wild-type (PspFI ThermoFisher Scientific cat. FD2224) or SID allele (BseGI, ThermoFisher Scientific cat. FD0874), respectively. Heterozygotes were also crossed with WT C57BL/6J to maintain the line.

All mice used in this study were bred and maintained at the University of Edinburgh animal facilities under standard conditions, and procedures were carried out by staff licensed by the UK Home Office under successive project licences (PPL numbers P3C15137E and PP4326006) and in accordance with the Animal and Scientific Procedures Act 1986 following initial approval by a local Animal Welfare and Ethical Review Board. All mice were housed within a SPF facility. They were maintained on a 12h light/dark cycle with an ambient temperature of 20–24 °C and relative humidity of 45–65%. They were housed in individually ventilated cages with wood chippings, tissue bedding and additional environmental enrichment in groups of up to five animals with *ad libitum* access to food and water. Mutant mice were caged with their wild-type littermates. After birth, heterozygous NTmut and SID animals were indistinguishable from wild-type in either sex. The sex of embryos (at embryonic days E3.5 and E9.5) was not determined post-mortem.

Plotting and statistical analysis of genotyping data were performed using the software GraphPad Prism version 10.5.0. The distribution of NT and SID genotypes was compared to expected Mendelian ratios using the Fisher’s exact test.

#### Cell culture

ESCs were grown in incubators at 37 °C with 5% CO_2_ in gelatin-coated tissue culture dishes. The culture medium was composed of Glasgow minimum essential medium (GMEM; Gibco cat. 11710035) supplemented with 15% foetal bovine serum (batch tested), 1x L-glutamine (Gibco cat. 25030024), 1x MEM non-essential amino acids (Gibco cat. 11140035), 1 mM sodium pyruvate (Gibco cat. 11360039), 0.1 mM 2-mercaptoethanol (Gibco cat. 31350010) and 100 U/ml leukaemia inhibitory factor (ESGRO® LIF, Merck cat. ESG1107, batch tested). For imaging, cells were directly cultured on gelatin-coated polymer coverslips (iBidi cat. 81156 or 80286). To rapidly degrade endogenous SALL4 in SALL4-dTAG ESCs, 500nM of dTAG-13 (Tocris cat. 6605) was added to the culture medium and cells were collected at the desired time for further analysis. For Western blot or TT-seq, cells were directly lysed on the plate. For timecourse CUT&RUN and ATAC-seq analyses, cells were cryopreserved in a mix of 40% culture medium (see composition above), 50% foetal bovine serum and 10% DMSO, and stored at -80°C until further processing.

*Sall4* knockout ^26^ (S4KO) and *MBD3* knockout (MBD3KO) ^58^ cell lines were kindly provided by Dr Brian Hendrich (University of Cambridge, UK). All cell lines generated in this study were derived from wild-type E14Ju09 ESCs, a clonal cell line derived from E14Tg2a ^84^. Cells were genetically engineered using the CRISPR/Cas9 technology, and following standard procedures ^85^. Briefly, cells were transiently transfected with mammalian expression plasmids coding for sgRNAs of interest (see Oligonucleotides section) and Cas9 fused to an EGFP fluorescent reporter (PX458, Addgene cat. 48138) using Lipofectamine 3000 reagents (ThermoFisher Scientific cat. L3000008) and following manufacturer’s protocol. When specific mutations were desired, a repair template for homologous recombination was co-transfected together with CRISPR reagents, either as a single-stranded DNA oligonucleotide (ssDNA for NTmut, SID, WTrec) or as a plasmid targeting vector (for SALL4-dTAG). Two days after transfection, Cas9-expressing cells were sorted by flow cytometry into gelatin-coated 96-well plates (1 cell/well) containing culture medium. Clones were expanded for 10-14 days and passaged into duplicate plates (one for freezing, one for genotyping). Genomic DNA from each clone was extracted from cell pellets using the DNeasy Blood & Tissue Kit (Qiagen cat. 69504) and following manufacturer’s protocol. Genotyping was performed by PCR amplification of targeted alleles (see primers below). Clones were screened by RFLP analysis, using specific restriction enzymes for NTmut allele (AlwNI, New England Biolabs cat. R0514S), SID allele (BseGI, ThermoFisher Scientific cat. FD0874) and WTrec allele (AseI, New England Biolabs cat. R0526S). For NTΔ and SALL4-dTAG alleles, clones were screened based on the observation of PCR bands of the correct size compared to wild-type genomic DNA. For all alleles, sequence identity was verified by Sanger sequencing of PCR products. This allowed us to confirm the homozygous/heterozygous status of each clone, and also to confirm that mutagenized sequences remained in frame with the rest of SALL4 coding sequence.

| **Cell lines** |  |  |  |
| --- | --- | --- | --- |
| **Name** | **Species** | **Cell type** | **Source / Reference** |
| Wild-type E14Ju09 | Mouse (*Mus musculus*) | Embryonic stem cells | Hooper et al., 1987 |
| Sall4 knockout (S4KO) | Mouse (*Mus musculus*) | Embryonic stem cells | Provided by Brian Hendrich laboratory (Miller et al., 2016) |
| MBD3 knockout (MBD3KO) | Mouse (*Mus musculus*) | Embryonic stem cells | Provided by Brian Hendrich laboratory (Kaji et al., 2006) |
| SALL4-dTAG | Mouse (*Mus musculus*) | Embryonic stem cells | This study |
| SALL4 NTmut | Mouse (*Mus musculus*) | Embryonic stem cells | This study |
| SALL4 NTΔ | Mouse (*Mus musculus*) | Embryonic stem cells | This study |
| SALL4 SID | Mouse (Mus musculus) | Embryonic stem cells | This study |
| SALL4 WTrec | Mouse (Mus musculus) | Embryonic stem cells | This study |

#### Immunofluorescence

Immunofluorescence was performed as described in a previous publication ^13^. Cells were washed with PBS and fixed with 4% PFA for 10min at room temperature. After fixation, cells were washed with PBS and permeabilised for 10min at room temperature in PBS supplemented with 0.3% (v/v) Triton X-100. Samples were blocked for 1h30min at room temperature in blocking buffer: PBS supplemented with 0.1% (v/v) Triton X-100, 1% (w/v) BSA and 3% (v/v) serum of the same species as secondary antibodies were raised in (ordered from Sigma-Aldrich). Following blocking, samples were incubated overnight at 4°C with primary antibodies (see Table below) diluted at the appropriate concentration in blocking buffer. After 4 washes in PBS supplemented with 0.1% (v/v) Triton X-100, samples were incubated for 2h at room temperature (in the dark) with fluorescently labelled secondary antibodies (Invitrogen Alexa Fluor Plus antibodies) diluted 1:500 in blocking buffer. Cells were washed 4 times with PBS supplemented with 0.1% (v/v) Triton X-100. DNA was stained with 1 µg/ml 4’,6-diamidino-2-phenylindole (DAPI) for 5min at room temperature, and finally mounted on coverslips using the ProLong Glass mounting medium (ThermoFisher Scientific cat. P36980). Images were acquired using the Zeiss LSM 880 confocal microscope with Airyscan module and the software Zeiss ZEN software (black edition). Images were subsequently processed and analysed using the software Fiji (based on ImageJ v1.54f).

#### Immunoprecipitation coupled with Western blot analysis

Co-immunoprecipitation and Western blot analyses were performed as described previously ^13^. Cells were washed with PBS, trypsinised and collected in 1.5ml Protein LoBind tubes (Eppendorf cat. 0030108116). To prepare nuclear protein extracts, cells were lysed for 20min on ice in 1ml of lysis buffer (10mM NaCl, 1mM MgCl_2_, 20mM HEPES pH7.5, 0.1% (v/v) Triton X-100) freshly supplemented with 1x protease inhibitor cocktail (Roche cat. 11873580001) and 0.5mM DTT. Nuclei were collected by centrifugation at 4°C for 10min at 500xg, and were resuspended in 1ml of lysis buffer freshly supplemented with 1x protease inhibitor cocktail, 0.5mM DTT and 250U of Benzonase nuclease (Merck cat. E1014-25KU). After a 5min incubation at room temperature, samples were supplemented with NaCl to obtain a final concentration of 150mM NaCl, and further incubated for 30min at 4°C on a rotating wheel. Tubes were centrifuged at 4°C for 30min at 16,000xg and supernatants (nuclear protein extracts) were transferred into new 1.5ml Protein LoBind tubes. Nuclear extracts were used directly for immunoprecipitation, or stored at -80°C. For input material, 50μl of protein extract was collected into a new tube, mixed with 1x volume of 2x Laemmli buffer (Sigma-Aldrich cat. S3401) and boiled for 5min at 90°C. Of note, for direct Western blot analysis (e.g. timecourse analysis of SALL4 expression in SALL4-dTAG ESCs), cells were directly lysed on the plate in Laemmli buffer and boiled for 5min at 90°C.

For immunoprecipitations, 5μg of antibody (see antibody list below) was added to each nuclear protein extract. Samples were incubated overnight at 4°C on a rotating wheel. 30μl of nProteinA Sepharose beads (Cytiva cat. 17528001), previously blocked with 0.5mg/ml BSA, were added to each nuclear extract and samples were incubated for 2h at 4°C on a rotating wheel. Samples were washed 5 times in lysis buffer freshly supplemented with 0.5mM DTT and 150mM NaCl. Between each wash, samples were centrifuged at 4°C for 1min at 500xg. After the final wash, immunoprecipitated material was eluted by boiling the beads for 5min at 90°C in 50µl of 2x Laemmli buffer.

Western blotting was performed according to standard procedures using the Mini-PROTEAN TGX system (Bio-Rad). Proteins were separated by SDS-PAGE and subsequently transferred onto a nitrocellulose membrane. The membrane was blocked for 1h at room temperature in PBS supplemented with 10% non-fat skimmed milk and 0.1% (v/v) Tween-20. The membrane was then incubated for 1h30min at room temperature with primary antibodies (see Table below) diluted at the appropriate concentration in PBS supplemented with 5% non-fat skimmed milk and 0.1% (v/v) Tween-20. The membrane was washed 4 times with PBS supplemented with 0.1% (v/v) Tween-20, and incubated for 2h at room temperature with fluorescently labelled secondary antibodies (Li-Cor IRDye) diluted in PBS supplemented with 5% non-fat skimmed milk and 0.1% (v/v) Tween-20. The membrane was finally washed 4 times with PBS supplemented with 0.1% (v/v) Tween-20. Western blot images were acquired using the Li-Cor Odyssey CLx system and analysed using the software Li-Cor Image Studio Lite version 5.2.5. The fluorescence signal measured within specific protein bands was normalised to a loading control (Histone H3) and plotted using the software GraphPad Prism version 10.5.0.

#### HDAC activity assay

Both nuclear protein extraction and SALL4 immunoprecipitation (from 50µg of protein extract, as measured by Bradford assay) were performed as described above. A fraction of the material was used for Western blot analysis (see relevant section) to evaluate the amount of SALL4 proteins in each immunoprecipitate. The histone deacetylase activity associated with SALL4 immunoprecipitates was quantified by chemiluminescence using the HDAC-Glo I/II Assay (Promega cat. G6420) according to manufacturer’s instructions. Quality control was performed to verify that quantified signal was specific (control treatment HDAC inhibitor Trichostatin A, control IP in *Sall4* knockout ESCs) and within the linear range of the assay (serial dilutions of a representative sample). For each immunoprecipitation, chemiluminescence measurements were performed in 3x technical replicates. Signal was normalised to the amount of SALL4 proteins in each immunoprecipitate (as measured by Western blot) and expressed relative to normalised HDAC activity in a wild-type control (SALL4 IP in E14Ju09 ESCs). Data was plotted using the software GraphPad Prism version 10.5.0.

#### Immunoprecipitation coupled with label-free mass spectrometry analysis (IP/MS)

Nuclear protein extracts were prepared as described above, but using buffers without any DTT. For immunoprecipitation, 50µg of antibodies were crosslinked onto 500µl of Protein G Dynabeads (ThermoFisher Scientific cat. 10004D) using 20mM DMP (ThermoFisher Scientific cat. 21666). 100µl of crosslinked beads (5µg of antibodies per IP) were mixed with each nuclear protein extract for 2h at 4°C on a rotating wheel. Samples were washed 3 times in wash buffer (150mM KCl, 50mM Tris-HCl pH8.0, 0.1% (v/v) Triton X-100). Between each wash, the supernatant was removed using a magnetic separator (ThermoFisher Scientific cat. 12321D). Immunoprecipitated material was eluted by incubating the beads in 50µl of 0.1% Rapigest (Waters cat. 186001861) for 15min at 60°C. The eluate was transferred into new 1.5ml Protein LoBind tubes and stored at -80°C before further processing.

Protein samples were reduced in 100mM DTT for 15min at 80°C and further processed by filter-aided sample preparation (FASP) ^86^ using commercial columns Vivacon 500 (Sartorius cat. VN01H21). We followed a standard procedure including removal of detergents (8M urea buffer), thiol alkylation with 100mM iodoacetamide (Merck cat. I1149) for 30min in the dark at room temperature and protein digestion with 0.6µg Trypsin (ThermoFisher Scientific cat. 90057) overnight at 37°C. Peptides were eluted from the column by centrifugation for 15min at 10,000xg. Following acidification (final concentration ≈0.4% trifluoroacetic acid), samples were loaded on a StageTip ^87,88^ for cleanup and concentration. Samples were eluted in buffer containing 80% acetonitrile and 0.1% trifluoroacetic acid. Following elution samples were dried and resuspended in 0.1% trifluoroacetic acid (5µl). Peptides (3µl) were separated on the UltiMate 3000 nano LC system fitted with 50cm EASY-Spray column using 190min gradient. MS data were acquired using the Orbitrap Fusion Lumos instrument operated in i.0n DDA mode. Parameters were as follows: cycle time was set to 3 sec, the MS1 scan Orbitrap resolution was set to 120,000, RF lens to 30%, AGC target was set to 4.0e5 and maximum injection time to 50ms. The MS2 scan was performed with the Ion Trap using rapid scan setting.

#### CUT&RUN

CUT&RUN was performed using a commercial kit (EpiCypher cat. 14-1048) and following the manufacturer’s protocol starting from 5 x10^5^ ESCs. For SALL4 CUT&RUN, freshly collected wild-type ESC (E14Ju09) were used. For timecourse analysis of H3K27ac in SALL4-dTAG ESCs, cryopreserved cells were thawed on the day of the experiment (see cell culture section). Cells were immobilised on Concanavalin A-coated beads, permeabilised with 0.01% digitonin and incubated with 0.5µg of desired antibodies (see antibodies list) overnight at 4°C on a nutator. Following washes, Protein A-Protein G-MNase fusion protein was added to the mix and activated by addition of 2mM CaCl_2_. Targeted DNA sequences were cleaved off by incubation for 1h at 4°C on a nutator. The reaction was quenched and supernatant containing fragmented genomic DNA was collected using a magnetic separator. DNA was subsequently cleaned up and concentrated by column purification. CUT&RUN libraries were prepared from ≈5ng of purified DNA using the KAPA Hyperprep Kit (Roche cat. 07962347001) together with KAPA dual-indexed adapters (Roche cat. 08278555702), with minor modifications favouring the preferential retention of small DNA fragments ^38^. CUT&RUN libraries were pooled in equimolar amounts and sequenced using the Illumina NovaSeq platform (Novogene Europe, UK).

#### Transient-transcriptome sequencing (TT-seq)

TT-seq in SALL4-dTAG ESCs was performed as described in a published protocol ^43^. Confluent 10cm dishes (≈10 x10^6^ cells) were treated for 2h with 500nM dTAG-13 (or DMSO as a control) to degrade endogenous SALL4, followed by a 15min incubation with 500µM 4SU to label nascent RNAs. Cells were directly lysed on the plate with 1ml of TRIzol (ThermoFisher Scientific cat. 15596026), the material was transferred into 1.5ml tubes and stored at -80°C. A replicate plate was used to check SALL4 protein levels by Western blot (quality control for dTAG treatment, see section above). Total RNA was extracted from samples using the Direct-zol RNA Miniprep Plus kit (Zymo, cat. R2070), following manufacturer’s protocol. 60µg of RNA was fragmented by treatment with 166.67mM NaOH for 20min on ice. 4SU-labelled RNA was biotinylated with the MTSEA biotin-XX linker (Biotium cat. 90066), and subsequently pulled down using the μMACS Streptavidin Kit (Miltenyi cat. 130-091-287). Following stringent washes, nascent RNA was eluted and concentrated using the RNeasy MinElute Cleanup Kit (Qiagen cat. 74204). 20 to 50ng of material was used for library preparation using the KAPA RNA Hyperprep Kit (Roche cat. 08098093702) together with dual-indexed adapters, using the protocol for degraded RNA samples. TT-seq libraries were pooled in equimolar amounts and sequenced using the Illumina NovaSeq platform (Novogene Europe, UK).

#### RNA-seq (ribosomal RNA-depleted libraries)

RNA-seq in *Sall4* mutant ESCs (NTmut, NTΔ, SID, WTrec, E14Ju09 control) was performed as described previously ^13^. All cell lines were seeded at the same density in 6-well plates, and collected after two days of culture for total RNA extraction using the RNeasy Plus Mini kit (Qiagen cat. 74134), following manufacturer’s instructions. Ribosomal RNA-depleted RNA-seq libraries were prepared using the KAPA RNA Hyperprep Kit (Roche cat. 08098131702) together with dual-indexed adapters, following manufacturer’s instructions. RNA-seq libraries were pooled in equimolar amounts and sequenced using the Illumina NovaSeq platform (Novogene Europe, UK).

#### ATAC-seq

ATAC-seq was performed by the company Azenta Life Sciences (South Plainfield, NJ, USA) from cryopreserved cells (see cell culture section). Cells were thawed, washed, and treated with DNAse I to remove genomic DNA contamination. Following dead cells removal via annexin bead conjugation, samples were quantified and assessed for viability using the Nexcelom Cellaca counter with AO/PI staining. After cell lysis and cytosol removal, nuclei were treated with Tn5 enzyme (Illumina cat. 20034197) for 30 minutes at 37°C. Tagmented DNA was purified using the MinElute PCR Purification Kit (Qiagen cat. 28004), barcoded using the Nextera XT Index Kit v2 (Illumina cat. FC-131-2001) and finally amplified by PCR. Libraries were cleaned up using SPRI beads, pooled in equimolar amounts and sequenced using the Illumina NovaSeq platform.

#### Antibodies

| Target | Host | Applications | Reference |
| --- | --- | --- | --- |
| SALL4 | Mouse (monoclonal) | Western blot, Immunoprecipitation, Immunofluorescence, CUT&RUN | Santa Cruz cat. sc-101147 |
| SALL4 | Rabbit (polyclonal) | Western blot, Immunofluorescence | Abcam cat. ab29112 |
| SALL1 | Mouse (monoclonal) | Western blot | Abcam cat. ab41974 |
| MTA1 | Mouse (monoclonal) | Western blot | Santa Cruz cat. sc-373765 |
| SIN3A | Rabbit (polyclonal) | Western blot | Abcam cat. ab3479 |
| MBD3 | Rabbit (monoclonal) | Immunoprecipitation | Abcam cat. ab240144 |
| OCT4 | Mouse (monoclonal) | Immunofluorescence | Santa Cruz cat. sc-5279 |
| H3K27ac | Rabbit (polyclonal) | CUT&RUN | Abcam cat. ab4729 |
| Histone H3 | Rabbit (polyclonal) | Western blot (loading control) | Abcam cat. ab1791 |
| Normal IgG (no target) | Rabbit (polyclonal) | CUT&RUN (negative control) | Epicypther cat. 13-0042 |

#### Oligonucleotides

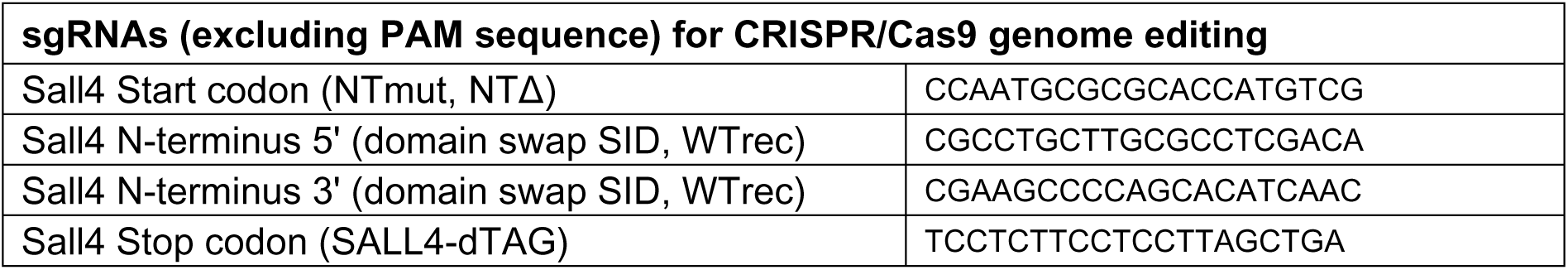
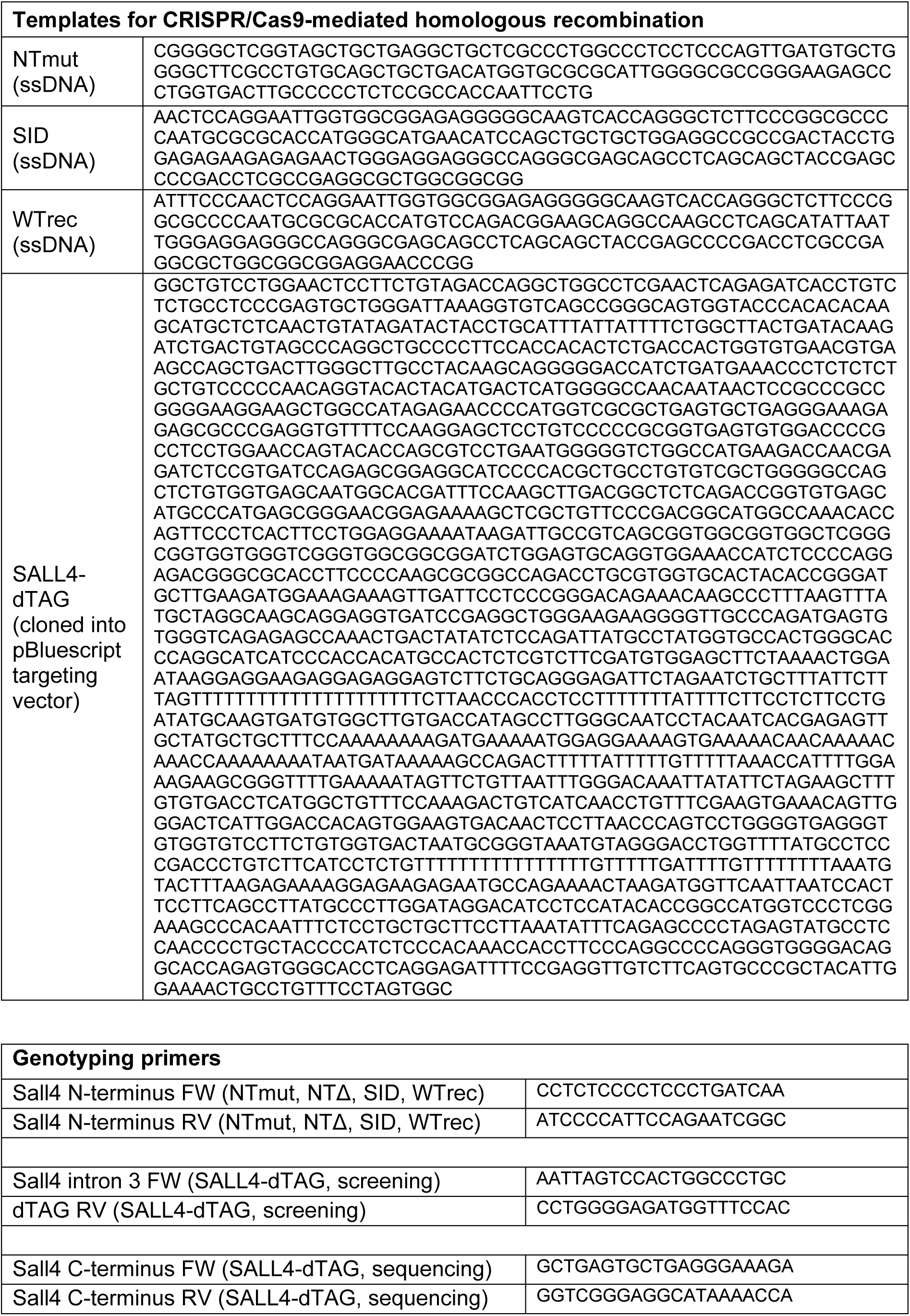

#### Bioinformatic and statistical analyses

##### CUT&RUN analysis

Paired-end sequencing reads were aligned to the mouse reference genome (mm10) to generate coordinate-sorted alignment files with Bowtie2 (v2.5.1). For each condition, biological replicates were merged to produce consolidated alignment files, which were indexed and summarised for alignment statistics. To visualise genome-wide signal, coverage tracks were generated from the merged BAM files. Properly paired fragments shorter than 1kb were retained, blacklisted regions were excluded, and coverage was calculated across the genome. Tracks were normalised using RPKM yielding bigWig files that were directly comparable across conditions. Peaks of enrichment were identified with MACS2 (v2.2.7.1) by comparing treatment samples against IgG controls, using paired-end mode and allowing all duplicate fragments to be retained, as appropriate for CUT&RUN data.

Genome-wide binding was assessed by partitioning the mouse genome into non-overlapping 10kb windows and computing log2-transformed RPKM values from CUT&RUN. For each window, the AT fraction was calculated and windows were ranked by AT content (40–80%). To visualise enrichment relative to AT richness, per-genotype signals were normalised to S4KO (log2 ratio to S4KO) and displayed as smoothed profiles (rolling median over 5,000 windows, sampled every 2,000). Shaded bands reflect local variability (rolling quantile range from 0.375 to 0.625). In addition, we repeated both analyses after explicitly removing reads overlapping called peaks and re-generating RPKM tracks, to evaluate whether observed global trends were driven solely by reads from the so-called peaks.

##### ATAC-seq analysis

For read alignment, the same strategy was performed as CUT&RUN analysis. Peaks of accessible chromatin were identified with MACS2 (v2.2.7.1) in paired-end mode. A consensus peak set was generated by merging peaks across all samples. Read counts within these peaks were obtained using *featureCounts*, and differential accessibility analysis was performed with *DESeq2* by comparing each dTAG-13 treatment timepoint against the DMSO control. Peaks with adjusted p-values below the significance threshold were considered differentially accessible.

##### Peak overlap and feature distribution analysis

To assess the co-localisation of binding and chromatin features, SALL4 CUT&RUN peaks were intersected with the ATAC-seq consensus peak set and H3K27ac CUT&RUN peaks using intervene tool. The differentially accessible peak set (describe in the above paragraph) from the 0.5h timepoint was subsequently intersected with the SALL4 CUT&RUN peak set to quantify co-occupancy. To assess proximity to target genes, differentially accessible peaks (0.5h) were overlapped with SALL4 target gene units (defined by TT-seq, from TSS minus 1kb to TES). A peak was considered associated if it fell within the defined target gene unit.

##### Transcriptomics data analysis

Reads were first aligned to a reference genome of mouse rDNA to remove ribosomal RNA contamination using Bowtie2 (v2.5.1) followed by alignment to mouse reference genome (mm10) with the STAR aligner (v2.7.10b). Post-alignment, the featureCounts tool (v2.0.6) was employed to quantify gene expression levels and differential gene expression analysis was performed using standard DESeq2 (v1.42.0) workflows, with adjusted p-values used to determine statistical significance (expression in mutant lines versus WT). For nascent transcription datasets (TT-seq), read counts were calculated across the whole gene (including introns) as opposed to exons only for steady-state transcription datasets (RNA-seq). To calculate the base composition of the gene unit, bedtools nuc command was used to quantify the AT content from the start and end coordinates of the gene entry in annotation (Gencode M25). Genes were grouped into equal-sized AT-content bins, and within each bin the median fold change was plotted. Error bands were defined as the 0.375–0.625 quantiles, representing the central 25% of values. Kendall’s tau correlations were calculated across all genes, and regression slopes (βbins) were derived from binned medians.

##### Identification of nascent transcription units

Nascent transcription units (TUs) were identified using hidden Markov models ^42^. To classify genomic windows, we overlapped the 10 kb windows with the detected TUs. Windows with at least 90% overlap with a TU were designated as *transcribed windows*. In contrast, windows that did not overlap by even a single base pair with any TU or with known gene annotations were designated as *non-transcribed windows*. This categorisation allowed separation of genomic regions into actively transcribed and non-transcribed classes for downstream chromatin analyses.

##### Correlation with SALL4 binding in 10 kb windows

The genome was partitioned into non-overlapping 10 kb windows, and SALL4 occupancy was quantified from CUT&RUN in wild-type cells. For each window, ATAC-seq and H3K27ac CUT&RUN changes were calculated as log2 ratios of dTAG-13 to DMSO RPKM for each timepoint. Windows were ranked by SALL4 occupancy, and profiles were plotted using the median values with error bands defined by the 0.375–0.625 quantiles (25% range). Kendall’s tau values were calculated for all windows, and regression slopes (βbins) were derived from binned means.

##### Explainable machine learning

To perform SHAP-based model interpretation, we first processed chromatin profiling data comprising publicly available ChIP-seq datasets and in-house CUT&RUN data. Raw reads for ChIP-seq experiments and their corresponding input controls were downloaded from GEO, whereas IgG CUT&RUN datasets were used as input controls for in-house data. All reads were processed using a standardized pipeline: alignment to the mm10 genome was performed using Bowtie2 (v2.5.1), followed by duplicate removal using samtools markdup. Signal tracks were generated using MACS2 (v2.2.9) to compute fold enrichment, which were exported as bedGraph files and subsequently converted to bigWig format. For each chromatin feature, summed signal intensities were calculated across gene promoters (±1 kb from TSS) and gene bodies (+1kb from TSS to TES) using bigWigAverageOverBed.

To assess feature importance in predicting RNA polymerase II occupancy, we trained a simple feedforward neural network implemented in PyTorch using five different train-validation-test data splits. Each model was optimized independently using a distinct set of hyperparameters selected through optimization with the Optuna framework. The following hyperparameters were tuned: L2 regularization strength (alpha, ranging from 1e-5 to 1e-1 on a log scale), number of neurons in the first and second hidden layers (hidden_layer_sizes_1 and hidden_layer_sizes_2, ranging from 16 to 64), and activation function (activation, chosen from “tanh”, “relu”, or “logistic”). Following model training, SHAP values were computed using DeepExplainer for each input feature (e.g., chromatin signals at promoter and gene body regions) on the test set. The background dataset for SHAP was generated using shap.kmeans, excluding any direct SALL4 target loci that overlapped with the training set to prevent information leakage. To account for variability in model training and to obtain robust estimates of feature importance, we independently trained and interpreted five models—one per data split—with different hyperparameters, allowing estimation of the variance in SHAP explanations across model configurations.

For comparative analysis of SHAP values, genes in the test sets were stratified based on statistical significance from a TT-seq differential expression experiment following acute SALL4 depletion, using adjusted p-value thresholds of 0.05, 0.5, 0.9, and 1.0. To further assess variability in SHAP values among subsets of significant genes, we randomly sampled 50 genes from each significance threshold group and computed the mean and median SHAP values across 50 iterations. This resampling approach allowed us to evaluate the stability of feature importance within specific transcriptional response categories.

##### Proteomic analysis

Raw proteomics data was analysed using the software MaxQuant ^89^ version 1.6.0.2. Data was further processed using the software Perseus 2.1.3.0 ^90^ with a standard workflow including sample annotation, filtering (reverse, identified by site, contaminants), logarithmic transformation and imputation of missing values. For each immunoprecipitation, quantified proteins were compared to a negative control (immunoprecipitation in *S4KO* or *MBD3KO* cell line, depending on the antibody used) using a Student’s t-test. Relative stoichiometries were determined as previously described ^59^ by dividing label-free quantification of proteins (iBAQ values) by the amount of immunoprecipitated proteins in each respective sample (SALL4 or MBD3 iBAQ, depending on the antibody used). Data was plotted using the software GraphPad Prism version 10.5.0.

#### Protein alignments

Proteins carrying a NuRD- or Sin3-interaction domain were identified by performing a BLAST search on the UniProt database ^91^ (https://www.uniprot.org/blast) covering the entire mouse proteome. The consensus sequence used for the NuRD-interaction domain was ‘MSRRKQAKPQHINWE’ (SALL4 residues 1-15) ^28,31^. The consensus sequence used for the Sin3-interaction domain was ‘GMNIQLLLEAADYLERRE’ (MXD1 residues 6-23) ^70^. Protein sequences corresponding to minimal NuRD- or Sin3-interaction domains were aligned using the Clustal Omega programme (https://www.uniprot.org/align) ^92^. Alignment files were uploaded on the ESPript platform (https://espript.ibcp.fr/ESPript/ESPript) ^93^ for visualisation.

### Data availability

Raw and processed data used to generate the figure panels (microscopy and Western blot images, datasheets underlying the plots, etc.) will be deposited on a public database at the time of publication.

High throughput sequencing data generated in this study is summarised in the table below.

|  |  |
| --- | --- |
| E-MTAB-16139 | TT-seq analysis of nascent transcripts following acute SALL4 degradation in mouse embryonic stem cells |
| E-MTAB-16160 | RNA-seq comparison of wild-type and N-terminal mutant (NTmut, SID) mouse embryonic stem cells |
| E-MTAB-16166 | Time-resolved H3K27ac CUT&RUN analysis following SALL4 degradation in mouse embryonic stem cells |
| E-MTAB-16168 | SALL4 CUT&RUN analysis in mouse embryonic stem cells |
| E-MTAB-16171 | Time-resolved ATAC-seq analysis following SALL4 degradation in mouse embryonic stem cells |

In addition to Table S1, other previously published datasets that were also used for analyses are described in the table below.

|  |  |
| --- | --- |
| GSM2417088 | H3K4me1 ChIP-seq analysis in mouse embryonic stem cells |
| GSM2417096 | H3K27ac ChIP-seq analysis in mouse embryonic stem cells |

Model weights and SHAP values from computational modelling will be available on the Github repository: https://github.com/kashyapchhatbar/SALL4-manuscript-2025.

Further information and requests for resources and reagents should be directed to Adrian Bird or Raphaël Pantier.

## Acknowledgements

We thank Rob Klose (University of Oxford) and Brian Hendrich (University of Cambridge) for sharing laboratory reagents and Verdiana Steccanella and Megan Brown for technical assistance with genotyping of animals. We also thank Fiona Rossi and Claire Cryer (Institute for Regeneration and Repair, University of Edinburgh) for support with flow cytometry and Toni McHugh and David Kelly (Wellcome Discovery Research Platform for Hidden Cell Biology, University of Edinburgh) for support with microscopy. We thank Martha Koerner and the University of Edinburgh Central Transgenic Core (CTC) for generation of transgenic animals.

This work was funded by the European Research Council Advanced Grant EC 694295 Gen-Epix and the Wellcome Investigator Award #107930. Imaging and mass spectrometry facilities were supported by a Core Grant to the Wellcome Centre for Cell Biology (#203149) as well as the Wellcome Discovery Research Platform for Hidden Cell Biology (#226791).

## Author contributions

Conceptualization, A.B., R.P. and K.C.; Methodology, R.P., K.C., T.Q., S.G., J.S., J.G., T.A. and C.P.; Software, K.C.; Formal Analysis, K.C. and R.P.; Investigation, R.P., K.C., T.Q., S.G., B.A.H., J.S., J.G., T.A., T.M. and C.P.; Writing – Original Draft, R.P., K.C. and A.B.; Writing – Review & Editing, R.P., K.C. and A.B.; Supervision, A.B., G.S. and R.P.; Funding acquisition, A.B.

## Supplementary Figures

**Figure S1.**
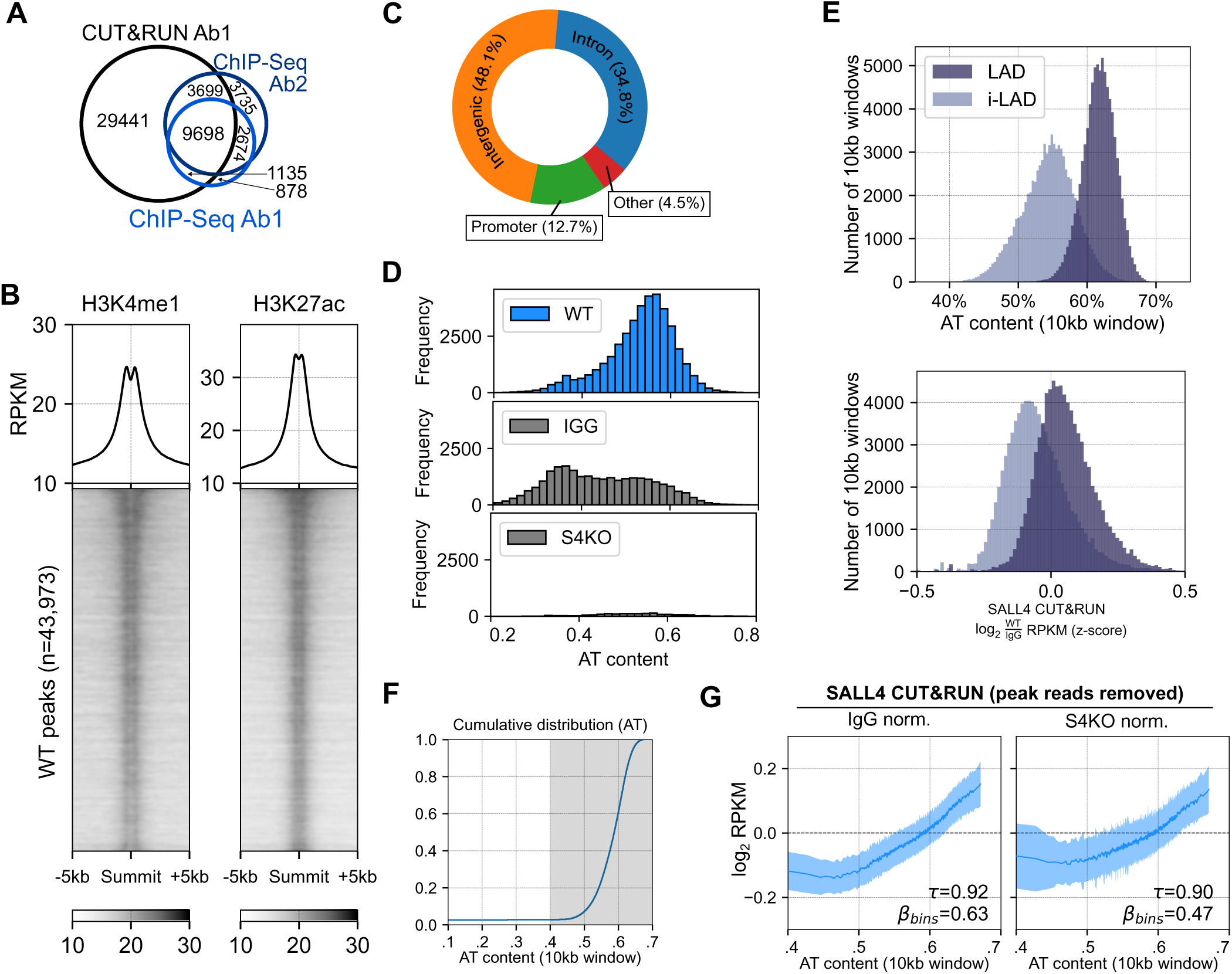
(related to Figure 1). **A.** Venn diagram showing the overlap of SALL4 CUT&RUN peaks with SALL4 ChIP-seq peaks detected in a previous study using two independent antibodies (Pantier et al, 2021). **B.** Heatmap showing SALL4 CUT&RUN, SALL4 ChIP-seq (Pantier et al, 2021), H3K27ac (Chronis et al., 2017) and H3K4me1 (Chronis et al., 2017) signal at SALL4 peaks detected in wild-type ESCs (n=43,973). **C.** Donut chart showing the genomic distribution of SALL4 CUT&RUN peaks. **D.** Bar graphs showing the distribution of SALL4 CUT&RUN peaks (blue) across varying levels of AT-content. Peaks detected in *Sall4* knockout ESCs or using normal IgG antibodies in wild-type cells, were used as negative controls (grey). **E.** Bar plot showing AT-content and normalised SALL4 CUT&RUN signal (z-score) within 10kb genomic windows in lamina-associated domains (LADs) or inter-LADs (iLADs). **F.** Cumulative plot showing the distribution of genomic bins from the mouse genome in relationship with AT-content. The shaded grey region highlights the AT-content where majority of the 10kb windows are accumulated. **G.** Plots showing normalised SALL4 CUT&RUN across 10kb genomic bins sorted by AT-content. SALL4 signal was calculated after removing reads within detected peaks (Fig. 1A) and was smoothed using a rolling mean with a window size of 1000 bins, and values were sampled every 400 bins. Shaded region represent one half standard deviation (SD) error across the 1000 bins.

**Figure S2.**
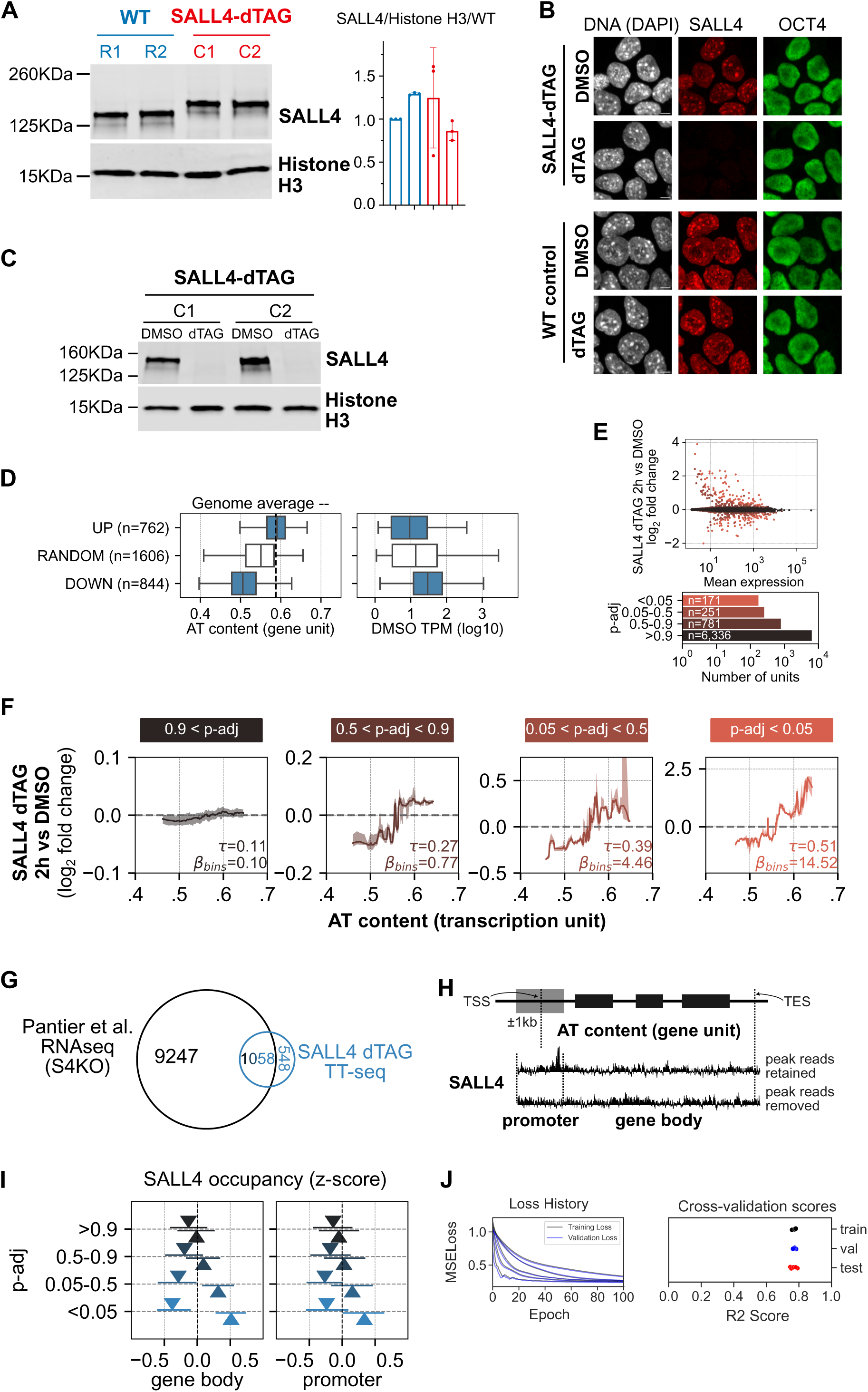
(related to Figure 2). **A.** Western blot analysis of SALL4 in SALL4-dTAG (untreated) compared to wild-type ESCs. Histone H3 was used as a loading control. Quantification is shown below the blots with each data point representing an independent replicate experiment (n=3). Error bars: Standard deviation (SD). **B.** Immunofluorescence analysis of SALL4 and OCT4 (pluripotency marker) in SALL4-dTAG and wild-type ESCs treated for 2h with dTAG-13, or DMSO as a control. DNA was stained with DAPI. Scale bars: 5µm. Images are representative from at least five randomly chosen fields of view for each condition, and data was reproduced in six independent SALL4-dTAG ESC clones. **C.** Western blot analysis of SALL4 in technical replicates of samples used for TT-seq (see Fig. 2a). Histone H3 was used as a loading control. **D.** Boxplots showing gene features of up-regulated (UP), down-regulated (DOWN), and randomly selected non-responsive (RANDOM) genes following acute SALL4 depletion. Left: AT content across the gene unit. Right: Steady-state expression levels in control (DMSO) cells, measured as log10 TPM. **E.** MA plot of nascent transcript changes following acute SALL4 degradation in ESCs (2 h dTAG-13 vs. DMSO). The accompanying bar plot shows numbers of differentially regulated transcription units stratified by adjusted p-value (padj) thresholds. Nascent transcripts were identified using an HMM-based approach ^42^ and overlapping annotated genes were removed prior to plotting. **F.** Gene expression changes (log₂ fold change) upon acute SALL4 depletion are plotted similarly across nascent transcripts detected from TT-seq data which do not overlap with known genes. Genes were grouped based on statistical significance and error bars represent standard deviation across each bin. The solid line represents the rolling median, and the shaded error band represents the inter-percentile range (0.375 to 0.625 quantiles). The strength of association is quantified using Kendall’s 𝜏 and the linear regression slope across the bins (β_bins_). **G.** Venn diagram showing the overlap of differentially expressed genes (p<0.05) detected by RNA-seq in mutant ESCs (Pantier et al, 2021) with direct transcriptional targets identified by nascent RNA-seq in degron cells. **H.** Visualisation of an arbitrary gene unit showing how promoter region (±1kb of TSS) and gene body (+1kb to TES) are defined and how SALL4 CUT&RUN signal across these regions is calculated. **I.** SALL4 CUT&RUN signal (IGG-normalized z-score) at gene bodies (left) and promtoers (right) is calculated after artifically removing peak reads across gene groups in Fig. 2C. Genes are further grouped into up-regulated (up arrow triangle) and down-regulated (down arrow triangle). The plotted triangles correspond to the median SALL4 occupancy and error bars represent the inter-percentile range (0.375 to 0.625 quantiles). **J.** Model performance for predicting RNA Pol-II occupancy from SALL4 CUT&RUN, H3K27ac ChIP-seq, and ATAC-seq (Table S2). Coefficients of determination (R^2^) are shown for training, validation and test datasets for 5 splits. Mean squared error (MSE) loss history for training (black) and validation (blue) over 100 training epochs, showing stable convergence of the model.

**Figure S3.**
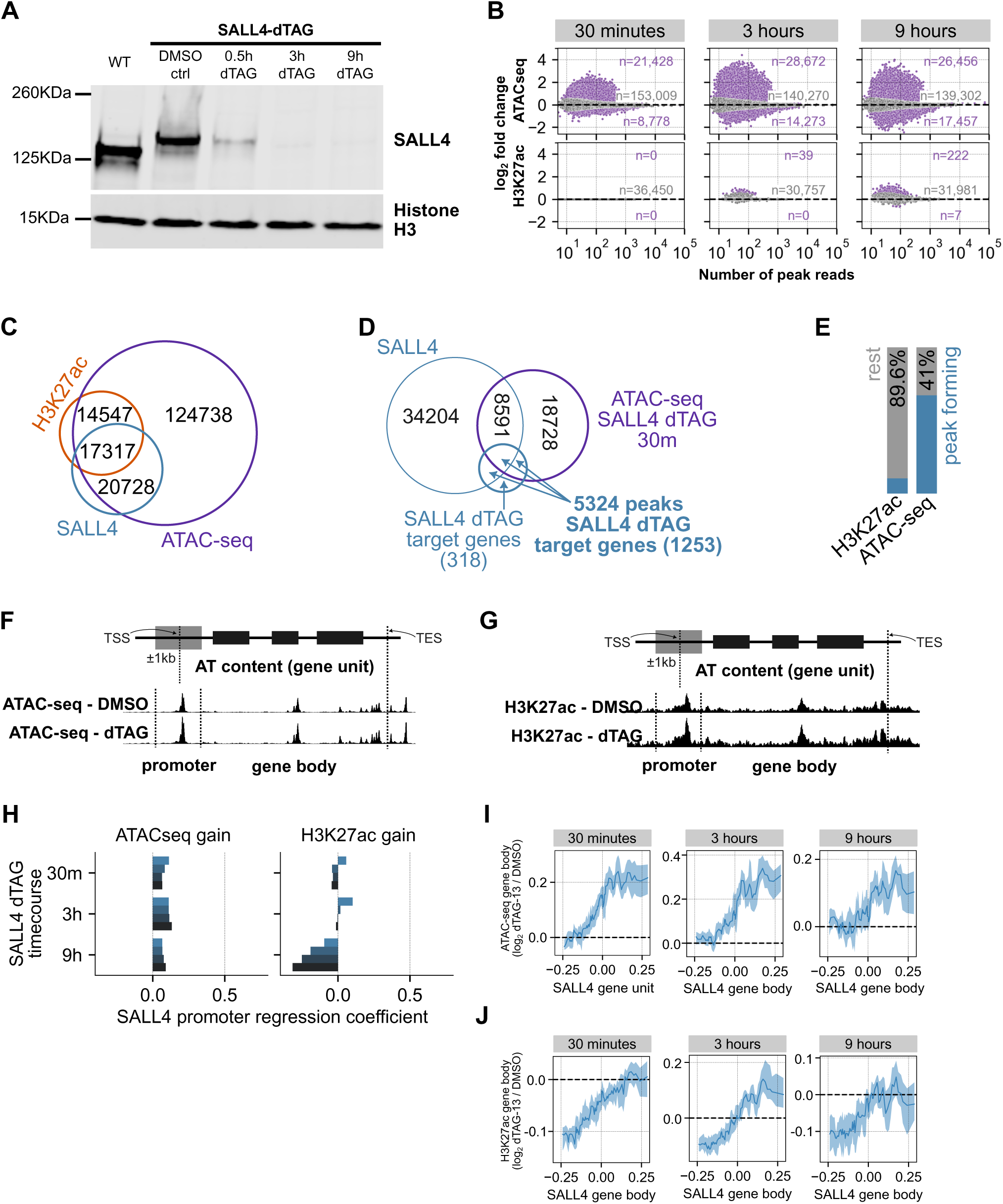
(related to Figure 3). **A.** Western blot analysis of SALL4 in technical replicates of samples used for ATAC-seq and CUT&RUN (see Fig.3A). Histone H3 was used as a loading control. Schematic of a representative gene unit showing **B.** MA plots showing the number of differentially accessible (top) and differentially acetylated (bottom) peaks (up- and down-regulated) after SALL4 degradation at 30 minutes, 3 hours, and 9 hours. **C.** Peak overlap between SALL4 CUT&RUN peaks, H3K27ac CUT&RUN peaks, and ATAC-seq peaks. **D.** Overlap of SALL4 peaks, differentially accessible ATAC-seq peaks 30 minutes after SALL4 degradation and gene bodies (1kb upstream of TSS to TES) of direct SALL4 targets (differentially regulated genes p-adj < 0.05 in Fig. 2C) **E.** Bar plot showing the proportion of H3K27ac CUT&RUN reads and ATAC-seq reads mapping to the mouse genome, either within (blue) or outside (grey) peaks. Visualisation of an arbitrary gene unit showing how promoter region (±1kb of TSS) and gene body (+1kb to TES) are defined and how **F.** ATAC-seq signal and **G.** H3K27ac signal across these regions is calculated. **H.** Bar plots of linear regression slopes across the bins (β_bins_) when promoter gain of ATAC-seq (left) or H3K27ac (right) is regressed against SALL4 occupancy. The colours represent the set of differentially regulated SALL4 target gene categories in Fig. 2C. Rolling median plots visualising changes in **I.** ATAC-seq and **J.** H3K27ac at SALL4 target gene bodies (p-adj < 0.05) at 0.5h, 3h and 9h post SALL4 degradation with respect to SALL4 CUT&RUN occupancy in WT mESCs. The shaded band represents the inter-percentile range (0.375 to 0.625 quantiles).

**Figure S4.**
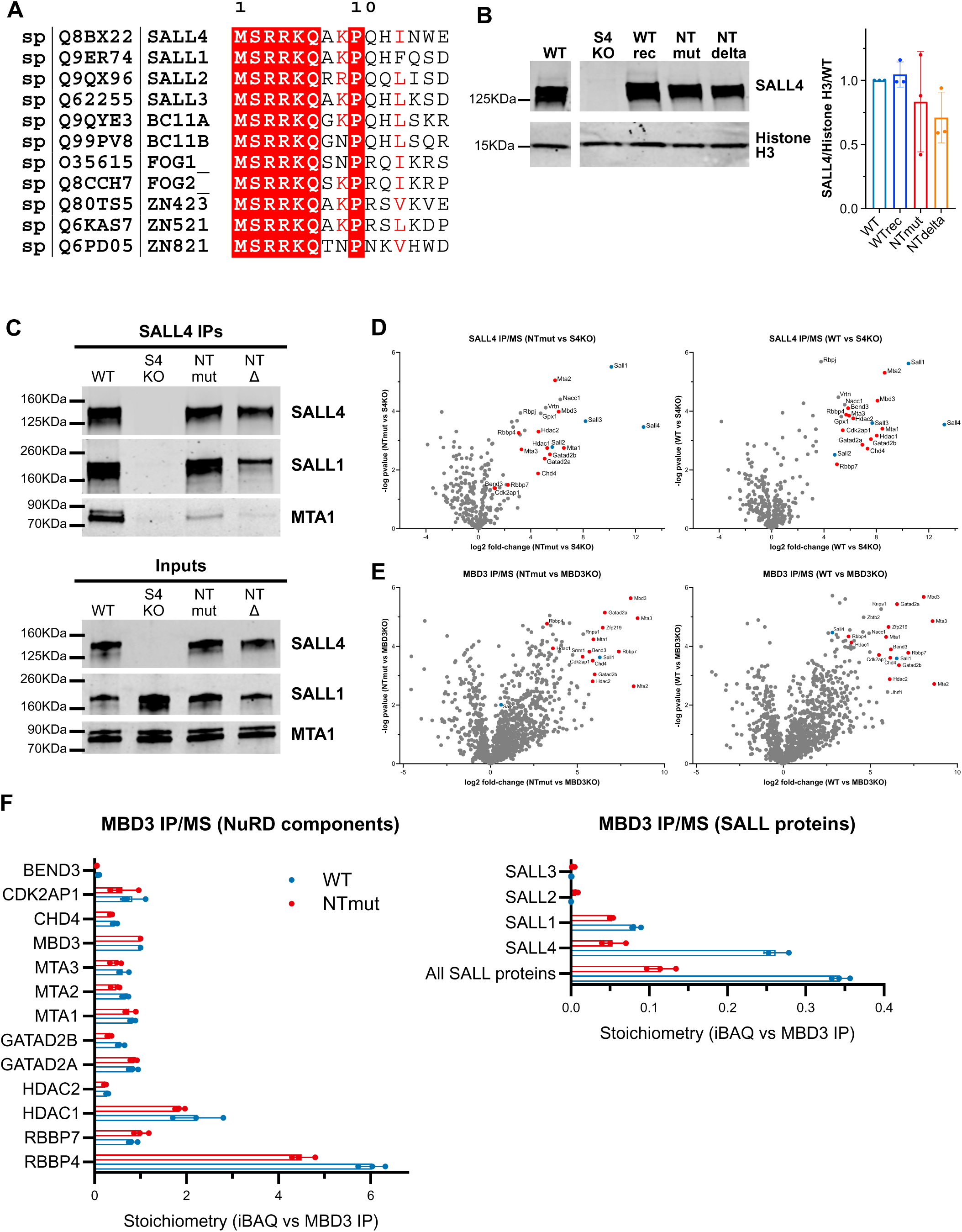
(related to Figure 4). **A.** Alignment of the NuRD-interaction of SALL4 with other mouse proteins containing a similar consensus sequence. Identical residues are shown in white on a red background; conservative substitutions are in red. **B.** Western blot analysis of SALL4 in mutant lines (WTrec, NTmut, NTΔ) compared to wild-type ESCs. Histone H3 was used as a loading control. For each staining, all samples shown are from the same Western blot membrane (non-adjacent lanes) with identical brightness/contrast ratios. Quantification is shown below the blots with each data point representing an independent replicate experiment (n=3). Error bars: Standard deviation (SD). **C.** Western blot analysis showing the co-immunoprecipitation of SALL4 with SALL1 (control) and MTA1 (NuRD subunit) in wild-type and N-terminal mutant ESCs. Images are representative from two independent replicate experiments. **D.** Vulcano plots showing the enrichment of proteins by SALL4 IP/MS in wild-type or NTmut ESCs compared to a negative control (*Sall4* knockout ESCs). NuRD subunits are highlighted in red and SALL proteins in blue. **E.** Vulcano plots showing the enrichment of proteins by MBD3 IP/MS in wild-type or NTmut ESCs compared to a negative control (*Mbd3* knockout ESCs). NuRD subunits are highlighted in red and SALL proteins in blue. **F.** Relative stoichiometries of NuRD subunits and SALL proteins following MBD3 IP/MS in wild-type or NTmut ESCs. Absolute protein amounts were estimated using iBAQ intensities and normalised to MBD3 IP in each sample. Each data point represents an independent replicate experiment (n=3). Error bars: Standard deviation (SD).

**Figure S5.**
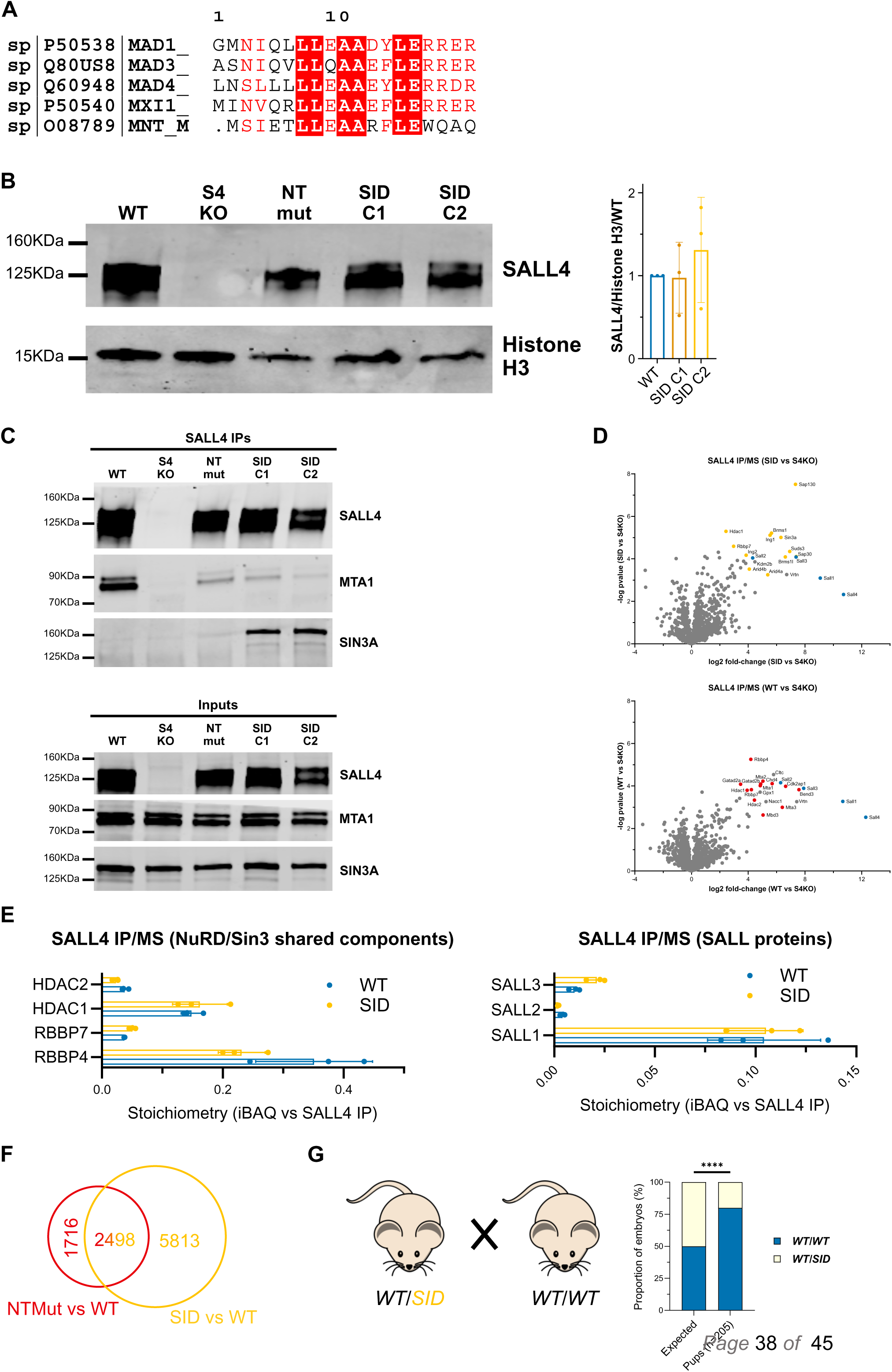
(related to Figure 5). **A.** Alignment of the Sin3-interaction of MAD1 with other mouse proteins containing a similar consensus sequence. Identical residues are shown in white on a red background; conservative substitutions are in red. **B.** Western blot analysis of SALL4 in mutant (SID) compared to wild-type ESCs. Histone H3 was used as a loading control. Quantification is shown below the blots with each data point representing an independent replicate experiment (n=3). Error bars: Standard deviation (SD). **C.** Western blot analysis showing the co-immunoprecipitation of SALL4 with SALL1 (control), MTA1 (NuRD subunit) and SIN3A (Sin3 subunit) in wild-type and N-terminal mutant ESCs. Images are representative from three independent replicate experiments. **D.** Vulcano plots showing the enrichment of proteins by SALL4 IP/MS in wild-type or SID ESCs compared to a negative control (*Sall4* knockout ESCs). NuRD subunits are highlighted in red, Sin3 subunits in yellow and SALL proteins in blue. **E.** Relative stoichiometries of shared NuRD/Sin3 subunits and SALL proteins following SALL4 IP/MS in wild-type or SID ESCs. Absolute protein amounts were estimated using iBAQ intensities and normalised to SALL4 IP in each sample. Each data point represents an independent replicate experiment (n=3). Error bars: Standard deviation (SD). **F.** Overlap of differentially regulated genes in NTMut/Δ ESCs and SID ESCs, as assessed by RNA-seq. **G.** Bar graphs showing the genotype of live pups (P21) following crosses between heterozygous *WT/SID* and wild-type mice. Statistical comparison with expected Mendelian ratios was performed using the Fisher’s exact test. Live pups show a strong bias towards a wild-type genotype, indicating that heterozygous SID mutation is embryonic lethal with partial penetrance.

## Supplementary Tables

Table S1. Gene Ontology (GO) term enrichment analysis of TT-seq up-regulated genes following acute degradation of SALL4.

Table S2. Gene Ontology (GO) term enrichment analysis of TT-seq down-regulated genes following acute degradation of SALL4.

**Table S3.**
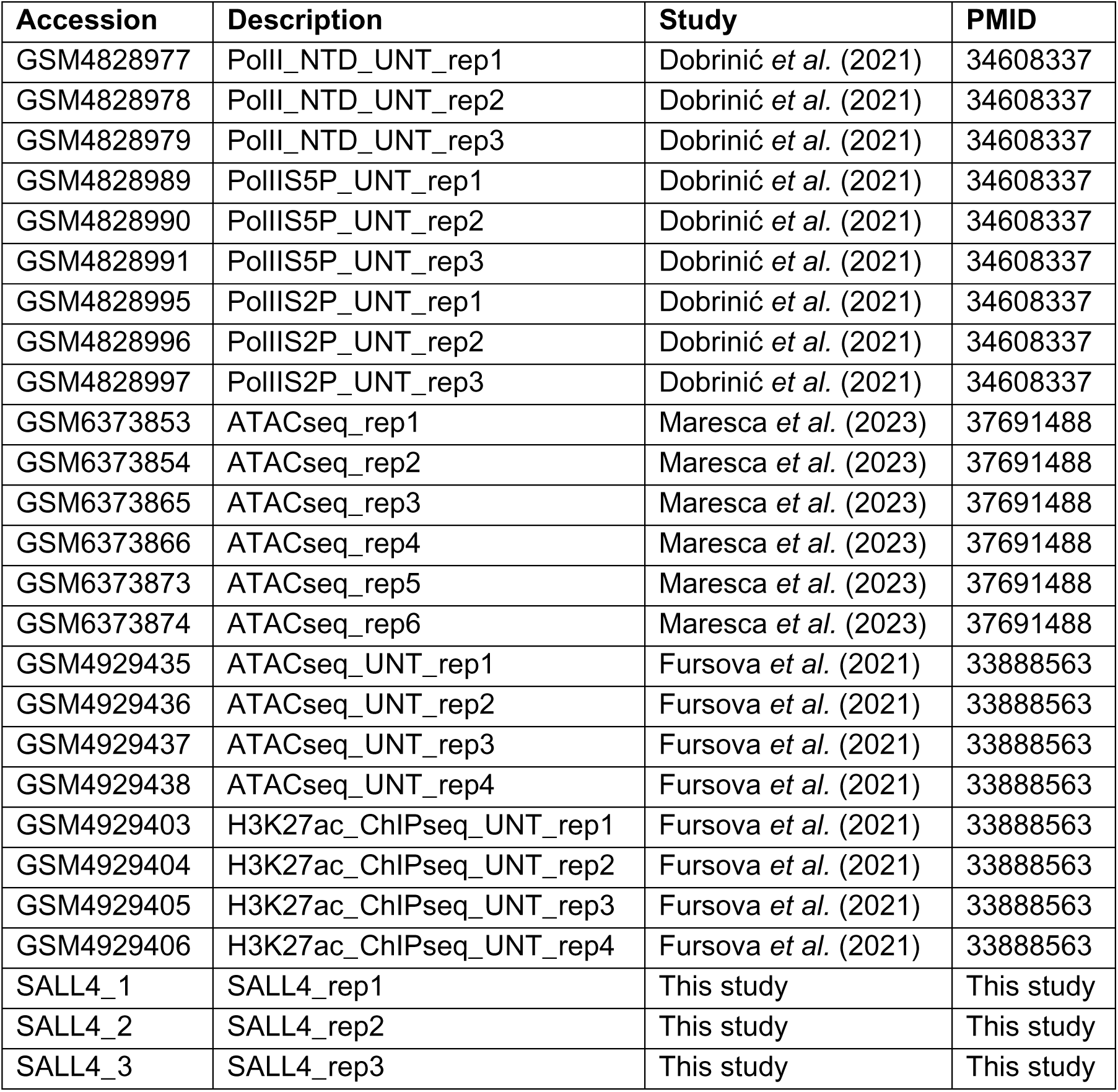
Public datasets and in-house profiles used as model inputs for RNA Pol II occupancy prediction in wild-type ESCs. GEO accessions, brief descriptions (replicates), source studies, and PMIDs are listed for: RNA Pol II (Dobrinić et al. 2021), ATAC-seq (Maresca et al. 2023; Fursova et al. 2021), H3K27ac ChIP-seq (Fursova et al. 2021), and SALL4 CUT&RUN (this study). For modelling, replicates were aggregated for each feature and total signal over promoters and gene bodies were quantified to create per-feature inputs.

Table S4. Gene Ontology (GO) term enrichment analysis of RNA-seq up-regulated genes in NTmut/Δ mouse embryonic stem cells.

Table S5. Gene Ontology (GO) term enrichment analysis of RNA-seq down-regulated genes in NTmut/Δ mouse embryonic stem cells.

Table S6. Gene Ontology (GO) term enrichment analysis of RNA-seq up-regulated genes in SID mouse embryonic stem cells.

Table S7. Gene Ontology (GO) term enrichment analysis of RNA-seq down-regulated genes in SID mouse embryonic stem cells.

For Tables S1, S2, S4, S5, S6 and S7, the analysis was conducted using the R package clusterProfiler. The table lists the most significantly enriched terms for the Biological Process (BP), Molecular Function (MF), and Cellular Component (CC) ontologies. These tables are attached as supplementary excel sheets.

